# The human visual cortex response to melanopsin-directed stimulation is accompanied by a distinct perceptual experience

**DOI:** 10.1101/138768

**Authors:** Manuel Spitschan, Andrew S. Bock, Jack Ryan, Giulia Frazzetta, David H. Brainard, Geoffrey K. Aguirre

**Author notes:** Contributions: M.S., D.H.B., and G.K.A. conceived the project. M.S. and G.K.A. designed the fMRI experiments. J.R., D.H.B., and G.K.A designed the perceptual experiment. M.S. and D.H.B. designed the spectral modulations. M.S., A.S.B., J.R., G.F. and G.K.A. collected fMRI data. G.F. collected pupillometry data. J.R. collected perceptual data. M.S., A.S.B. and G.F. analyzed fMRI data. M.S. and G.F. analyzed pupillometry data. G.K.A. implemented temporal models for the fMRI and pupillometry data. D.H.B., J.R. and G.K.A. analyzed perceptual data. M.S. analyzed the effects of biological variability upon photoreceptor contrast. G.K.A. created the figures. M.S. and G.K.A. wrote the manuscript with contributions from J.R., A.S.B., G.F. and D.H.B.

## Abstract

The photopigment melanopsin supports reflexive visual functions in people, such as pupil constriction and circadian photoentrainment. What contribution melanopsin makes to conscious visual perception is less studied. We devised a stimulus that targeted melanopsin separately from the cones using pulsed (3 s) spectral modulations around a photopic background. Pupil-lometry confirmed that the melanopsin stimulus drives a retinal mechanism distinct from luminance. In each of four subjects, a functional MRI response in area V1 was found. This response scaled with melanopic contrast and was not easily explained by imprecision in the silencing of the cones. Twenty additional subjects then observed melanopsin pulses and provided a structured rating of the perceptual experience. Melanopsin stimulation was described as an unpleasant, blurry, minimal brightening that quickly faded. We conclude that isolated stimulation of melanopsin is likely associated with a response within the cortical visual pathway and with an evoked conscious percept.

## Introduction

Human visual perception under daylight conditions is well described by the combination of signals from the short (S)-, medium (M)-, and long (L)-wavelength cones.^1^ Melanopsin-containing, intrinsically photosensitive retinal ganglion cells (ipRGCs) are also active in bright light (Figure 1a). The ipRGCs have notably prolonged responses to changes in light level, and thus signal retinal irradiance in their tonic firing.^2^ Studies in rodents, non-human primates, and people have emphasized the role of the ipRGCs in reflexive, non-image forming visual functions that integrate information over tens of seconds to hours, such as circadian photoentrainment, pupil control, and somatosensory discomfort from bright light.^3–6^

**Figure 1:**
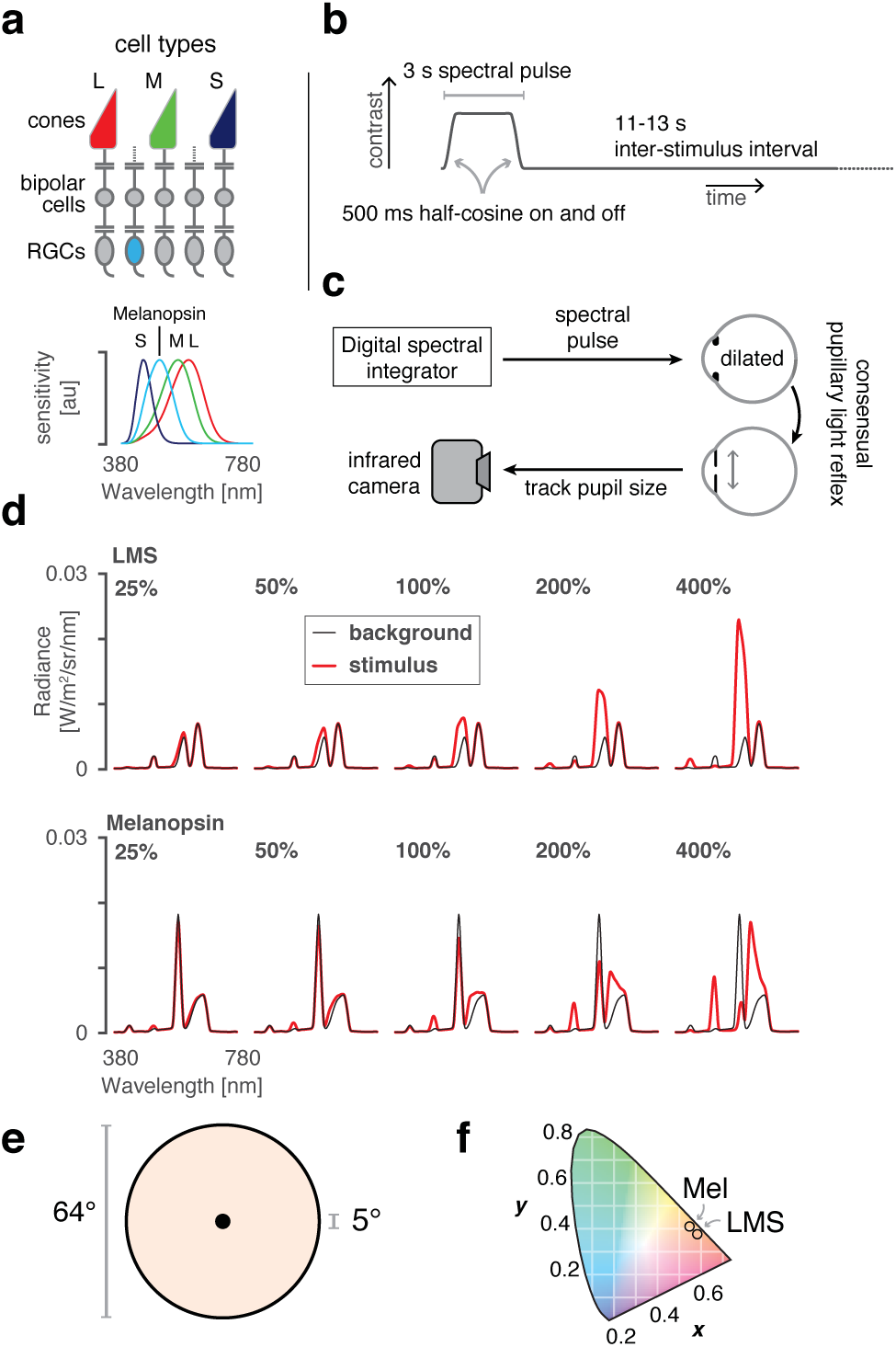
Overview and experimental design. **(a)** *Top* The L, M, and S cones, and melanopsin-containing ipRGCs, mediate visual function at daytime light levels. *Bottom* The spectral sensitivities of these photoreceptor classes. **(b)** Multiple 3-second, pulsed spectral modulations were presented, windowed by a 500 ms half-cosine at onset and offset, and followed by an 11-13 s ISI. A given experiment presented either a single contrast level, or multiple contrast levels in a counter-balanced order. **(c)** During fMRI scanning, subjects viewed pulsed spectral modulations, produced by a digital spectral integrator, with their pharmacologically dilated right eye. The consensual pupil response of the left eye was recorded in some experiments. **(d)** Stimulus spectra. Changes between a background spectrum (black) and modulation spectra (red) targeted a given photoreceptor channel with varying degrees of contrast. *Top* Spectra targeting the L, M, and S cones and thus the post-receptoral luminance channel. We use the terms “LMS” and “luminance” interchangeably to describe this stimulus. The nominal melanopic contrast for these modulations was zero. *Bottom* The corresponding spectra for stimuli targeting melanopsin. The nominal L-, M-, and S-cone contrast of these stimuli was zero. **(e)** Spectra were presented on a uniform field of 64°(visual angle) diameter. Subjects fixated the center of a 5° masked region, minimizing stimulation of the macula. **(f)** The calculated chromaticity of the background spectra was approximately matched for the LMS and melanopsin directed stimuli, and had a light-orange hue.

Relatively unexamined is the effect of melanopsin phototransduction upon visual perception, which operates at shorter timescales.^7–9^ In addition to tonic firing, ipRGCs exhibit transient responses to flashes of light with an onset latency as short as 200 ms.^10^ Several ipRGC subtypes project to the lateral geniculate nucleus, where they are found to drive both transient and tonic neural responses.^11^ As the geniculate is the starting point of the cortical pathway for visual perception, it is possible that ipRGC activity has an explicit visual perceptual correlate.

Here we examine whether isolated melanopsin stimulation drives responses within human visual cortex, and characterize the associated perceptual experience. Our approach uses tailored modulations of the spectral content of a light stimulus, allowing melanopsin to be targeted separately from the cones in visually normal subjects.^12,13^ We also studied the converse modulation, which drives the cone-based luminance channel while minimizing melanopsin stimulation. We collected blood oxygen level dependent (BOLD) functional magnetic resonance imaging (fMRI) data while subjects viewed brief (three-second) pulses of these spectral modulations. Concurrent infrared pupillometry was used to confirm that our stimuli elicit responses from distinct retinal mechanisms. Finally, we characterized the perceptual experience of selective melanopsin-directed stimulation, and examined whether this experience is distinct from that caused by stimulation of the cones.

## Results

Four subjects were studied in multiple experiments while they viewed intermittent pulses of spectral contrast directed at either the post-receptoral luminance pathway (LMS, equal contrast on cones) or the melanopsin containing ipRGCs (Figure 1a, 1b). During functional MRI scanning, subjects viewed these stimuli with their pharmacologically dilated right eye; in some experiments the consensual response of the left pupil was also recorded with an infra-red camera (Figure 1c). Different stimuli produced contrast upon the targeted photoreceptors between 25% and 400% (Figure 1d; additional stimulus details in Figure S1). The subject maintained fixation upon a masked central disk (Figure 1e), while spectral changes occurred in the visual periphery against a background that was depleted in short-wavelength light and thus had a light-orange hue (Figure 1f).

### V1 cortex responds to melanopsin contrast

We first examined the extent of cortical response to high-contrast spectral pulses. Each subject viewed approximately 200 pulses each of the 400% luminance and melanopsin stimuli. We measured the reliability of the evoked response within subject, and then at a second level across subjects and the two hemispheres. Pulses of luminance contrast that minimized melanopsin stimulation (Figure 2a) produced responses in the early cortical visual areas, generally corresponding to the retinotopic projection of the stimulated portion of the visual field.^14^ Spectral pulses directed at melanopsin that minimized cone stimulation also evoked responses within the visual cortex (Figure 2b). In subsequent experiments, we examined the evoked responses to luminance and melanopsin stimulation within a region of interest in V1 cortex that lies entirely within the retinotopic projection of the stimulated visual field. The time-series data and evoked responses from within this region for the initial, 400% contrast only experiment can be found in Figure S2.

**Figure 2:**
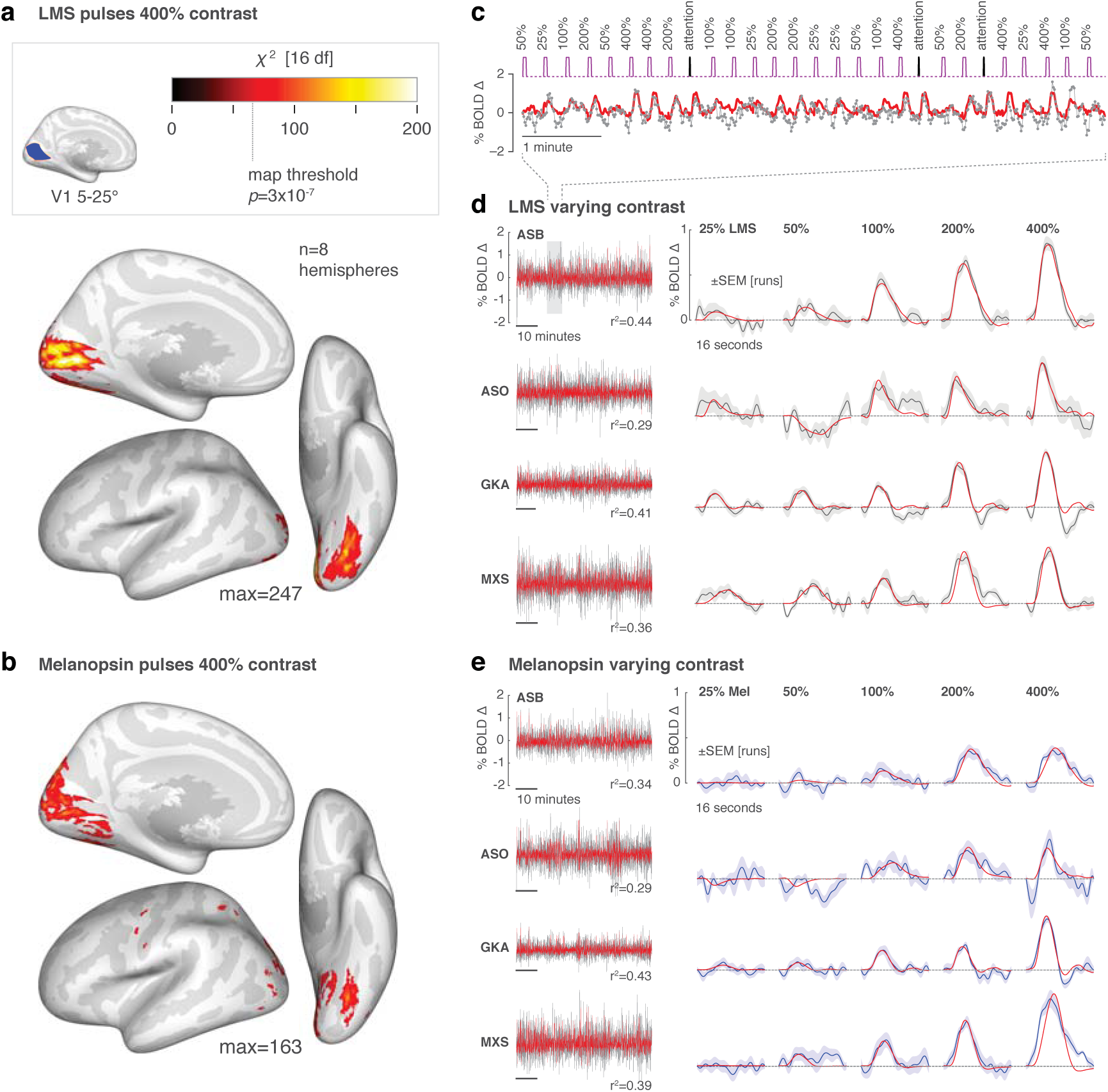
Visual cortex responses to LMS and melanopsin contrast. **(a)** Cortical response to pulses of 400% LMS contrast across subjects and hemispheres. Threshold corresponds to a map-wise *α* = 0.05 (Bonferroni corrected for the number of vertices). Inset *top* is the region of V1 cortex with retinotopic representation corresponding to the visual field range of 5-25 ° radial eccentricity, indicated in blue. Subsequent analyses examine the mean signal from this region. **(b)** The corresponding surface map obtained in response to 400% Melanopsin contrast pulses. **(c)** Example fit (red) of the Fourier basis set to a portion of the BOLD fMRI time-series data (gray). **(d)** V1 responses to LMS stimulation of varying contrast. *Left* The BOLD fMRI time-series data from the area V1 region for each subject (black), following pre-processing to remove nuisance effects. A Fourier basis set modeled (red) the mean evoked response to each contrast level with the *r*^2^ value of the model fit indicated. *Right* The evoked responses for each subject and stimulus level (black), and SEM of the response across the 9-11 scanning runs performed in each subject (shaded region). The responses were fit by a model (red) that convolved a step function of neural activity by the hemodynamic response function measured for each subject (Figure S3). **(e)** The corresponding responses within the V1 region to melanopsin stimulation of varying contrast.

If the visual cortex encodes information from the ipRGCs, we would expect that the degree of BOLD fMRI response should reflect variation in the degree of melanopsin stimulation, similar to the modulation of cortical response seen to variation in luminance contrast.^15^ Each of the four observers was studied again, this time with spectral pulses that varied in the degree of contrast upon the LMS or melanopsin channels. Figure 2c shows an example of the data obtained from the V1 region of interest in response to luminance pulses during one scan run for one observer. The time-series was fit with a Fourier basis set that estimated the shape of the BOLD fMRI response evoked by stimuli of each contrast level. Figure 2d presents the time-series data and evoked responses for the four subjects during luminance stimulation. Luminance pulses evoked consistent responses in the V1 region of interest, with a steadily increasing amplitude of evoked response across contrast levels. Variation in melanopic contrast (Figure 2e) produced similar data, with an increasing amplitude of BOLD fMRI response to larger contrasts.

We fit the evoked responses at each contrast level for each subject using an empirical measure of the subject’s hemodynamic response function, along with parameters that controlled the duration of an underlying neural response and the amplitude of the evoked BOLD fMRI signal (Figure S3). We obtained the amplitude of response as a function of contrast for each subject and each stimulus (Figure 3; LMS and melanopsin; grey and blue lines, respectively).

**Figure 3:**
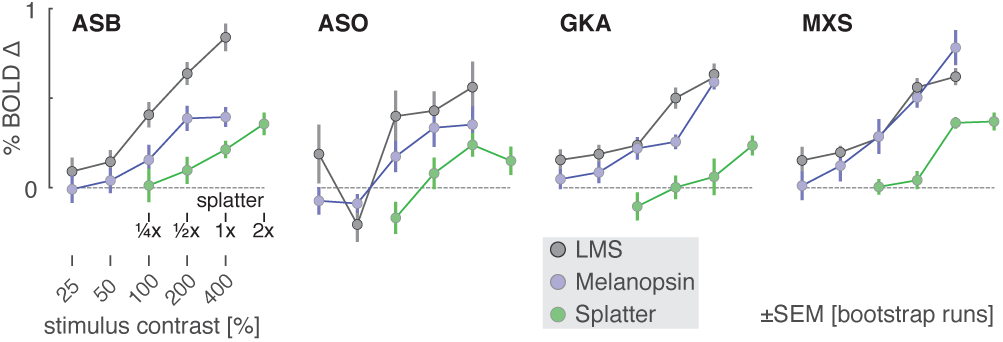
V1 BOLD fMRI response by stimulus contrast. The amplitude of evoked response with the V1 region was obtained for each subject and contrast level for the luminance (gray), melanopsin (cyan), and “splatter” (green) stimulus conditions. The 1x splatter condition presented cone contrast equal to the maximal inadvertent contrast (resulting from imperfections in device control) estimated from measurements of the spectra in the melanopsin experiments.

As suggested by the evoked responses in Figure 2, the measured amplitude increased as a function of contrast for both luminance and melanopsin stimulation for all four observers. While we modeled the duration of underlying neural activity, the results did not support the claim of a distinct temporal response to melanopsin stimulation (Figure S4).

While the melanopsin-directed spectral pulses were designed to produce no differential stimulation of the cones, biological variation and inevitable imperfection in device control results in some degree of unwanted cone stimulation (termed “splatter”).^12,13,16^ We considered the possibility that what appeared to be a visual cortex response to melanopsin contrast was in fact a response to the small amount of cone contrast inadvertently produced by our nominally cone silent spectral pulses.

We obtained spectroradiometric measurements of the stimuli that were actually produced by our device at the time of the BOLD fMRI experiment for each subject. For each of these measurements we calculated the inadvertent contrast that the cones would have experienced within these 400% melanopsin modulations in a biologically typical subject. We took the maximum contrast values calculated for the measurements across subjects, and created a new spectral pulse that was designed to have no melanopsin stimulation, but to have cone contrast equal to this estimate of inadvertent contrast. Scaled versions of this modulation corresponded to logarithmically-spaced larger (2x) and smaller 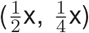 multiples of the “splatter” contrast. We again studied the four subjects with BOLD fMRI while they viewed these stimuli, and measured the amplitude of response as a function of splatter contrast (Figure 3, green line). In all four subjects, the melanopsin response function was larger than the splatter response function. This indicates that the cortical response to melanopsin cannot be explained entirely by imperfection in stimulus generation. We then explored if biological variability could result in a greater degree of inadvertent cone contrast than our analysis of device imprecision alone would suggest. Our characterization of the stimuli in terms of cone contrast relies upon assumed values for several biological variables, including lens density, peak spectral sensitivity of the cone photopigments, their density, and the density of macular pigment. We conducted simulations in which we calculated the degree of inadvertent cone contrast that would have resulted given deviations from our assumptions, following estimated distributions of these biological variables.^17^ We find that it is very unlikely (approximately one chance in 100,000) that the responses observed in the four subjects could have resulted solely from inadvertent cone contrast (Figure S5).

The spectral sensitivity of the rod photoreceptors overlaps extensively with that of melanopsin. The background used for our melanopsin-directed stimuli was 3.5 log_10_ scotopic Trolands (scot Td), nominally at or above the rod saturation threshold, found to be 3.0 log_10_ scot Td (Figure 2 of Adelson 1982)^18^ or 3.3-3.7 log_10_ scot Td (Aguilar & Stiles 1954).^19^ Therefore, we expect in our experiments that there is no, or minimal, time-varying signal contributed by the rods. We attempted in a control experiment to further exclude this possibility by making use of an assumed difference in temporal sensitivity of the rods and melanopsin, but this experiment was uninformative (Figure S6). We return to this topic in the discussion.

A prior functional MRI study that presented a 50% Weber contrast melanopsin modulation did not find responses within the visual cortex, but did observe BOLD fMRI responses within the frontal eye fields.^20^ The authors speculated that melanopsin stimulation produces changes in alertness that manifest as these cortical responses, although eye movements were not recorded during their study. In our whole brain analysis (Figure 2a, 2b) we find responses within the frontal eye fields for both the luminance and melanopsin pulses at lowered map thresholds (unthresholded maps available from http://neurovault.org/collections/2459/). We considered the possibility that our stimulus pulses might cause subjects to briefly increase or decrease saccadic eye movements. We measured variation in eye position during the 3 s of stimulation and during the interstimulus interval (Figure S7). Subjects consistently reduced eye movements during the luminance and melanopsin stimulation periods as compared to the inter-stimulus-interval. This effect may account for the frontal eye field responses in our data and in the prior report.^20^ As eye movements alone can evoke responses in visual cortex,^21^ we considered that a systematic difference in eye movements across contrast levels might confound our finding of a contrast-dependent response in area V1. However, no eye movement difference was seen as a function of contrast level or stimulus type (LMS vs. melanopsin).

### Different kinetics of pupil response to melanopic and luminance pulses

We have previously shown using sinusoidal spectral modulations that pupil responses to melanopsin stimulation have different temporal properties as compared to the responses evoked by modulations of luminance.^13^ In the current study, we recorded pupil responses to pulsed spectral modulations during the presentation of melanopsin and LMS stimulation of varying contrast. We examined these pupil responses for qualitative differences in the time course of the response. Such a demonstration would increase confidence that our stimuli target distinct retinal mechanisms.

The average pupil response was obtained for each contrast level and stimulus type. In the across-subject averages (Figure 4a; individual subject data in Figure S8), an evoked response to LMS stimulation is seen at even the lowest contrast level (25%). As LMS contrast grows, the evoked pupil response becomes larger, with distinct features corresponding to the onset and the offset of the 3 s stimulus pulse. The response to melanopsin contrast (Figure 4b) begins smaller, but also increases with contrast. Unlike the pupil response to LMS contrast, it is difficult to discern an indication of stimulus offset in the extended response to melanopsin stimulation.

**Figure 4:**
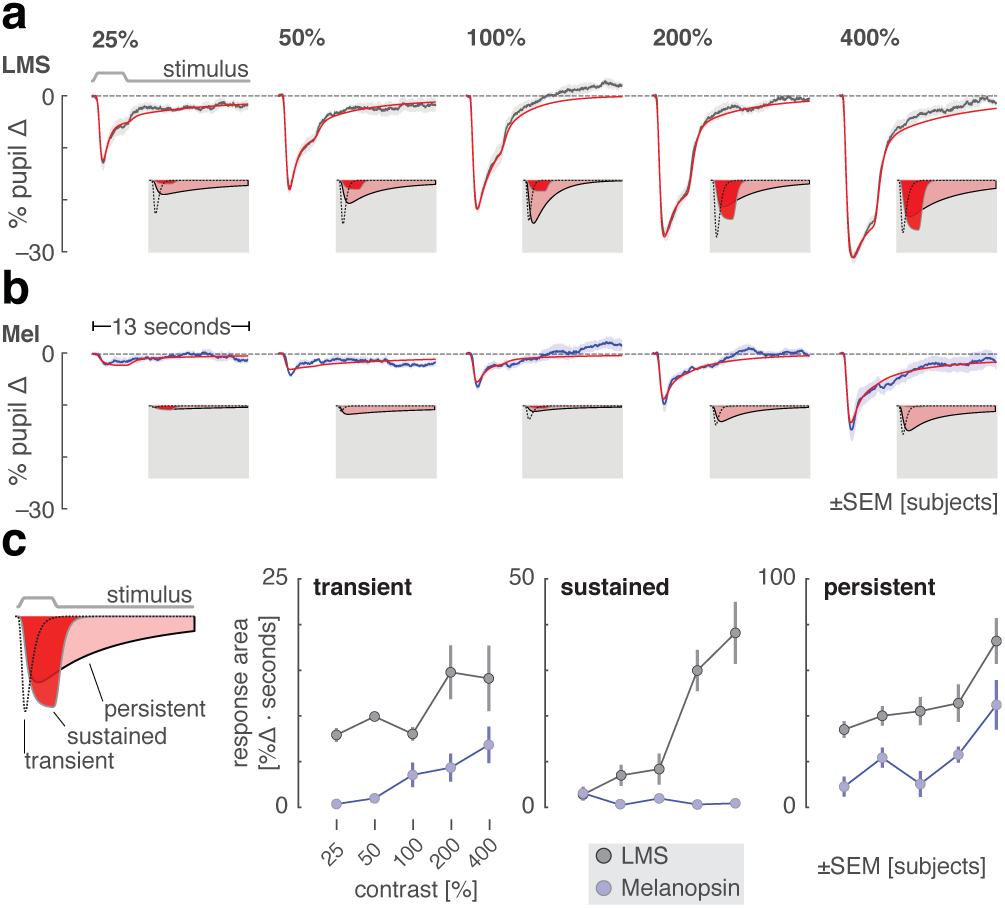
Consensual pupil responses to LMS and melanopsin stimulation. The consensual pupil response of the left eye was measured during stimulation of the pharmacologically dilated right eye. **(a)** The mean (across subjects) pupil response evoked by LMS stimulation of varying contrast levels (black), with SEM across subjects (shaded). The evoked response was fit with a three component, six-parameter model (red). The three components that model each response are shown inset on a gray field. **(b)** The corresponding mean pupil responses evoked by melanopsin stimulation of varying contrast levels. **(c)** Amplitude of the three model components as a function of stimulus contrast. Inset *left* is an illustration of the three model components. *Right* gain parameter for each model component as a function of contrast for LMS (gray) and melanopsin (blue) stimulation.

We quantified these observations by fitting a temporal model (Figure S9) to the average evoked pupil responses. The model has three temporally distinct components that capture an initial transient constriction of the pupil at stimulus onset, a sustained response that tracks the stimulus profile, and a persistent response as the pupil slowly re-dilates in the seconds after stimulus offset (shown inset in each plot panel in Figure 4a and 4b, and schematically inset left in Figure 4c). The amplitude of each of these components was measured as a function of contrast for the LMS and melanopsin stimuli (Figure 4c; temporal parameter values in Figure S10). The amplitude of both the initial transient and persistent response increase with LMS and melanopsin contrast. The behavior of the sustained component, however, is different for the two types of stimulation. Luminance contrast produces steadily increasing sustained pupil constriction that is time-locked to the profile of the stimulus. In contrast, there is essentially no component of this kind in the melanopsin-driven pupil response. This behavior is in keeping with the temporally low-pass properties of the melanopsin system.^13^ We verified that the qualitative difference between the pupil response to luminance and melanopsin contrast remained when an alternative fitting procedure that locked the temporal profile of each model component across stimulation conditions was employed.

### Melanopsin stimulation evokes a distinct visual percept

We find that a melanopsin-directed spectral pulse evokes a measurable response in the visual cortex. This suggests that people have conscious perceptual awareness of stimulation of the ipRGCs. Prior studies have found that melanopsin contrast contributes to a sensation of brightness, as subjects rate lights that contain melanopsin and luminance contrast as brighter than a light with luminance contrast alone.^7^ We were curious as to whether the perception of selective melanopsin-directed contrast appears simply as the typical experience of “brightness” conveyed by the luminance channel, or if there is a distinct perceptual experience associated with our melanopsin-directed stimulus.

We recruited 20 subjects and asked them to view 400% contrast pulses of LMS, melanopsin, and a stimulus changing in power by an equal multiplicative factor across all wavelengths, thus stimulating both melanopsin and luminance channels (“light flux”). Subjects were asked to rate nine perceptual qualities of the light pulse, each quality defined by a pair of antonyms (e.g., dim to bright). Subjects were not informed of the different identities of the stimuli, and the order was randomized as described in Online Methods. Subjects were also invited to offer their free-form observations at the end of the study during a debriefing session (summarized in Table S2).

A challenge of such measurements is the psychophysical sensitivity of the human visual system to even small amounts of differential cone contrast.^22,23^ We implemented additional stimulus calibration measures to further reduce spectral variation due to device instability (see Online Methods). In the measured stimulus spectra, the amount of inadvertent cone contrast in the melanopsin-directed stimulus due to imprecision in stimulus control was small (Figure S11).

Subjects rated each property of each stimulus twice, allowing us to confirm that within-subject reliability was high (across-subject mean Spearman correlation of test-retest reliability = 0.73 ± 0.18 SD). Additionally, there was good subject agreement in the ratings (across-subject mean Spearman correlation of ratings from one left-out subject to mean ratings of all other subjects = 0.53 ± 0.13 SD).

Subjects consistently rated the melanopsin stimulus as perceptually distinct from the LMS or light flux pulses (Table S1). We summarized these measurements by submitting them to a principal components analysis (Figure 5a). The first and second dimensions explained 35% and 19% of the variance in ratings, respectively. Within this space a support vector machine could classify subject responses to melanopsin as distinct from those for LMS or light flux with 92% cross-validated accuracy. A plot of the weights that define the classification dimension (Figure 5b) reveals the primary qualities of melanopsin stimulation. To these subjects, and in our own experience, the onset of the melanopsin contrast appears as a somewhat unpleasant, blurry, minimal brightening of the field. Most notably, however, this percept is fleeting, and rapidly followed by a fading or loss of perception from the stimulus field. Many of the subjects described the melanopsin stimulus pulse as being colored. This was typically with a yellow-orange appearance, although three subjects reported a greenish percept.

**Figure 5:**
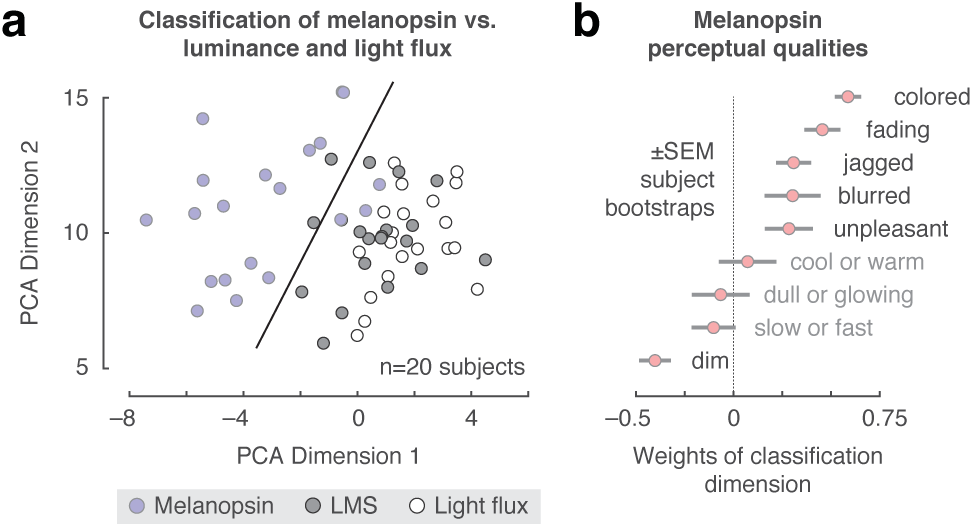
Perceptual ratings of melanopsin, luminance, and light flux. Subjects rated nine qualities of spectral pulses that targeted melanopsin, luminance, and their combination (light flux). **(a)** The set of perceptual ratings were subjected to a principal components analysis. Each point corresponds to the ratings provided by one subject for one stimulus type within the space defined by the first two dimensions of the PCA solution. A linear support-vector machine was trained to distinguish ratings for melanopsin stimulation from the other two stimulus types within this twodimensional space. The classification boundary is shown. **(b)** The classification dimension (normal to the classification boundary) describes how melanopsin stimulation was perceived differently from light flux and luminance. The mean weights (across boot-strap resamples) that define the classification dimension are shown.

The perceptual ratings of the LMS and light flux stimuli were quite similar, with the LMS rated as having more color (again perhaps due to the inadvertent chromatic contrast present in the stimulus; Figure S11) and the light flux as being brighter. Prior studies have found that melanopsin contrast is additive to LMS contrast in the perception of brightness.^7^ In our data, this would be consistent with higher ratings on the dim-to-bright scale for light flux pulses as compared to LMS. A post-hoc test supported this interpretation (Wilcoxon signed-rank test of dim-to-bright ratings in Light Flux compared to LMS: p=0.0088).

## Discussion

Our studies indicate a role for the melanopsin-containing ipRGCs in conscious human vision. We find that high-contrast spectral exchanges designed to isolate melanopsin evoke responses in human visual cortex. Pupil responses to these stimuli are distinct from those produced by luminance contrast, consistent with separation of retinal mechanisms. The cortical response is not easily explained by inadvertent stimulation of the cones and is associated with a distinct perceptual experience.

Previous studies in rodents and humans with outer photoreceptor defects have suggested that the visual cortex responds to melanopsin stimulation. Zaidi and colleagues reported the case of an 87 year-old woman with autosomal-dominant cone-rod dystrophy who was able to correctly report the presence of an intense, 480 nm 10 s light pulse, but not other wavelengths.^24^ Similarly, in mice lacking rods and cones, the presentation of a narrowband 447 nm light evoked a hemodynamic (optical imaging) signal change in the rodent visual cortex, with a slightly delayed onset (1 s) and a reduced amplitude as compared to the same measurement in a wild-type mouse.^25^ In our work we measured cortical and perceptual responses to melanopsin-directed stimulation in the intact human visual system.

### A cortical response

The melanopsin containing ipRGCs have broad projections to sub-cortical sites.^26^ Studies in the rodent and primate demonstrate as well projections to the lateral geniculate nucleus, where evoked responses to melanopsin stimulation can be found.^11,25,27^ Whether these signals are further transmitted to the visual cortex in normally sighted humans or non-human animals has been unknown. We find that pulsed melanopsin stimulation evokes contrast-graded responses within primary visual cortex. Responses to the highest (400%) contrast stimulus extend into adjacent, retinotopically organized visual areas, including ventrally in the vicinity of the peripheral representation for hV4 and VO1;^28^ a similar spatial distribution of cortical responses was observed to luminance stimulation.

By using a background depleted in short-wavelength light,^8^ we created substantial melanopic contrast in our stimuli, albeit ~3.5x less than is available in rodent models with a shifted long-wavelength cone.^27^ We found that 100% contrast pulses were required to obtain a measurable cortical response to melanopsin. The contrast response functions for both V1 fMRI amplitude and persistent pupil constriction appeared to be in the linear range and rising even at our maximum, 400% contrast level.

A characteristic property of the ipRGCs is their tonic firing to transient stimuli. Our model of the evoked BOLD responses in V1 estimates the underlying duration of neural activity (Figure S3). We observed an increasing duration of neural activity in response to melanopsin stimulation across contrast levels, which was not seen in response to luminance stimulation (Figure S4). We regard this result as tentative, however, principally because a similar, increasing duration of neural response was seen for the “splatter” control modulation.

### A visual percept

Consistent with the presence of a V1 neural response, we find that melanopsin-directed stimulation is accompanied by a distinct visual percept. We viewed these stimuli over many hours of experiments, and ourselves experienced the onset of the melanopsin spectral pulse as a diffuse, minimal brightening of the visual field. The appearance was curiously unpleasant.

The diffuse, even blurry, property of the percept might be related to the broad receptive fields of neurons driven by melanopsin stimulation,^29^ consistent with the extensive dendritic arbors of the ipRGCs.^30^ In a prior study, subjects reported that lights appear brighter when melanopsin contrast is added to the stimulation of the cone-based luminance channel.^7^ We find a conceptually similar effect in our data, as subjects rated pulses of light flux (which contain melanopic contrast) as brighter than pulses with cone contrast alone.

The most striking aspect of the percept evoked by the melanopsin pulse is that the brief brightening is then followed by a fading of perception of the stimulus field, on occasion spreading to involve the masked macular region of the stimulus. This was subjectively similar to Troxler fading. This aspect was remarked upon by several of our observers: “[the experience was] like blinding”; and “[the fade] to black that is the noise when your eyes are closed” or “kind of like if you got hit in the head really sharply … flashing lights and fade out.” (Table S2). The melanopsin containing ipRGCs send recurrent axon collaterals to the inner plexiform layer where they are positioned to modulate cone signals.^31^ Consistent with this, melanopic contrast has been shown to attenuate cone-driven electroretinogram responses in the rodent over minutes.^27^ The prominent and rapid experience of fading for our melanopsin-directed stimulus perhaps reflects the unopposed action of this attenuation mechanism.

Our data do not allow us to determine if one or more of the reported perceptual experiences arising from melanopsin stimulation are a direct consequence of ipRGC signals arriving at visual cortex sites, or from the interaction of melanopsin and cone signals at earlier points in the visual pathway.

### The challenge of photoreceptor isolation

Our conclusions depend upon the successful isolation of targeted photoreceptor channels. Measurements and simulations indicate that the functional MRI results are unlikely to be explained by inadvertent cone contrast from known sources of biological variation (Figure S5).^17^ Nonetheless, we think it prudent to carry forward concern regarding inadvertent cone intrusion, and to search for additional means to exclude this possible influence. For example, in the current study we examined in the functional MRI data whether there was a difference in the time-course of response to luminance and melanopsin-directed stimuli, but did not find convincing evidence of such (Figure S3). A time-course dissociation in the fMRI data would have provided further support—similar to that obtained in the pupil data—that our stimuli drive distinct mechanisms. Different temporal profiles of stimulation may afford greater traction on this question in future studies.

In our perceptual experiment, the melanopsin stimulus was reported to have a change in hue. This was usually, but not universally, reported as a yellow-orange. In this experiment we do not have available an estimate of the amount of reported color change that may be attributable to imperfections in cone silencing. Consequently, we are unable to reject the possibility that small amounts of chromatic splatter produce this percept.

Our results are also subject to any systematic deviation of photoreceptor sensitivity from that assumed in the design of our spectral modulations. One example model deviation is the presence of “penumbral” cones that lie in the shadow of blood vessels, and thus receive the stimulus spectrum after it has passed through the hemoglobin transmittance function. These photoreceptors can be inadvertently stimulated by a melanopsin-directed modulation, producing a percept of the retinal blood vessels when the spectra are rapidly flickered (≥ 4 Hz).^16^ While it is possible to also silence the penumbral cones in the melanopsin stimulus,^12^ this markedly reduces available contrast upon melanopsin (below 100%). We circumvented this problem here by windowing the onset of the melanopsin stimulus with a gradual transition (effectively 1 Hz) that removed the penumbral cone percept from our stimulus pulse.

We did not explicitly silence rods in our melanopsin-directed stimulus. Our background is at light levels considered to be above rod intrusion, and we have previously demonstrated a pupil response to melanopsin-directed modulation around a background an order of magnitude brighter,^13^ indicating that the melanopsin system responds at light levels well above rod intrusion. In principle, we could further exclude the possibility of rod intrusion by examining a flickering version of our melanopsin-directed stimulus. In such an experiment we would identify a flicker frequency at which rods could respond (if not saturated) but for which melanopsin might not be expected to do so (e.g., 4-8 Hz). Finding no cortical response to the stimulus would support the contention that the rods are saturated. In practice, this control experiment faces two challenges. First, melanopsin may still respond within this frequency range.^10^ Second, this stimulus may drive the penumbral cones, producing a percept of the blood vessels and a cortical response.^12,16^ Modifying the stimulus to silence the penumbral cones would markedly reduce available contrast on both the rods and melanopsin, defeating the purpose of the experiment. Nonetheless, we attempted this control study and obtained uninformative results (Figure S6). An important area for future investigation is the relationship between rod and melanopsin signals in the transition between mesopic and photopic vision.

We note that these challenges attend our prior study of cortical responses to rapid melanopsin flicker.^12^ In those experiments, penumbral-cone silent, sinusoidal melanopsin modulations with 16% Michelson contrast were studied. For comparison to the stimuli used in the current study, we can express contrast as the peak of the sinusoid relative to the trough. This yields ~40% Weber contrast. Given our finding here that roughly 100% Weber contrast was needed to evoke a V1 response, we now regard our prior study as not fully resolving the possibility that rapid modulation of the ipRGCs drives a cortical response.

The question of whether melanopsin contributes to visual perception at photopic light levels in people is one of considerable interest, as it challenges the orthodoxy that only three photopigments contribute to daylight vision. Two previous studies using silent substitution methodology reported psychophysical sensitivity in detection of cone-silent spectral modulations at photopic light levels.^8,9^ These studies also faced the challenge of photoreceptor isolation, as even small imperfections in the silencing of cones could lead to detection. An inferential strength of the current study is that we measure a graded, supra-threshold visual cortex response to varying contrast levels, which we may compare to the effect of imprecision in cone silencing. Further, presentation of supra-threshold contrast allows for the characterization of the appearance of the stimulus, as was done here.

## Conclusions

Our results suggest that people can “see” with melanopsin. The high-contrast, melanopsin-directed spectral modulation we studied is a distinctly unnatural stimulus, but a valuable tool for demonstrating the presence of a melanopic signal in the cortical visual pathway. Many of our subjects found the melanopsin-directed stimulus to be unpleasant to view. We are curious if variation in the perceptual or cortical response to this stimulus is related to the symptom of photophobia.^32^ Under naturalistic conditions, it appears that melanopsin adjusts the sensitivity of the cone pathways.^27^ The interaction of melanopsin and cone signals in human vision is an exciting avenue for investigation, particularly given recent findings of a role for melanopsin in the coarse spatial coding of light intensity.^29^

## Methods

A digital light synthesis engine (OneLight Spectra) was used to produce spectral modulations that targeted either the melanopsin photopigment or the LMS cones with varying contrast (25%, 50%, 100%, 200% and 400%) against a rod-saturating background (100-200 cd/m^2^; >3.3 log sc td). Pulse stimuli (3s, cosine windowed at onset and offset) were presented within a wide-field, uniform annulus with an outer diameter of 64° and an inner diameter of 5°, minimizing macular stimulation. Stimuli were adjusted for each observer’s nominal age to account for age-specific pre-receptoral filtering (see **Online Methods**, *Visual stimuli*). The quality of photopigment isolation was assessed by combining spectroradiometric measurements of the stimuli with a resampling approach that modeled sources of biological variation in photoreceptor spectral sensitivity (see **Online Methods**, *Simulation of biological variability causing inadvertent cone contrast*).

Four observers (four men; aged 27, 28, 32, 46; three of whom are authors of this study) viewed the stimuli with their pharmacologically dilated right eye while they underwent functional MRI in a 3T Siemens Prisma MRI scanner with a 64-channel headcoil. The consensual pupillary response to the stimuli was measured from the left eye during some scanning sessions using an infrared eye tracker. Stimulus pulses were jittered in their onset timing and spaced 14–16 seconds apart. Subjects were asked to detect an occasional, brief (500 msec) dimming of the stimulus field to which they made a button press. This served to monitor subject alertness and provided events that were used to derive a hemodynamic response function (HRF) for each observer.

BOLD fMRI data underwent standard pre-processing and were projected to a spherical atlas of cortical surface topology, supporting anatomical definition of the location and organization of retinotopic cortex (see **Online Methods**, *MRI data acquisition and initial processing*). Because stimuli were presented asynchronously with respect to fMRI acquisitions, the time-series data were fit with a Fourier basis set to extract the average evoked response to each stimulus type. The resulting evoked response per stimulus type was then fit with a two-parameter model incorporating the duration of an underlying step of neural activity, and the amplitude of this response after convolution by the subject-specific HRF (see **Online Methods**, *BOLD fMRI time-series analysis*).

In a separate experiment, conducted outside of the scanner, 20 observers (9 men, 11 women; mean age 27, range 20–33) viewed the LMS and melanopsin-directed stimuli, as well as pulses of broadband spectral change (light flux) which stimulated both cones and melanopsin. These observers were not involved in the design and conduct of the study and were not informed as to the identity of the pulses. They were asked to rate the stimuli along nine perceptual dimensions, given as antonym pairs (see **Online Methods**, *Perceptual rating experiment*).

The research was approved by the University of Pennsylvania Institutional Review Board and conducted in accordance with the principles of the Declaration of Helsinki. All subjects gave written informed consent. All experiments were pre-registered in the Open Science Framework. All data and code are available.

Detailed methods are described in **Online Methods**.

## Acknowledgements

This work was supported by the National Institutes of Health (Grant R01 EY024681 to G.K.A. and D.H.B., Core Grant for Vision Research P30 EY001583, and Neuroscience Neuroimaging Center Core Grant P30 NS045839), the Department of Defense (Grant MR141251 to G.K.A). We thank Fred Letterio for technical assistance, and Andrew S. Olsen for his assistance with data collection.

## Online Methods

### Pre-registration

The experiments were the subject of pre-registration documents. Data collection followed the pre-registration documents in regard to the number of subjects, extent of data collection, stimulus generation, and exclusion criteria. In some cases addenda were submitted to the preregistration before data collection began, with the pre-registered protocol being that which includes the modifications specified in these pre-data-collection addenda. In some cases the analysis approach presented in this paper differs from that described in the pre-registered protocol. Table S3 lists all pre-registration documents by experiment and deviations from the registered protocols. Some deviations were detailed in addenda submitted after data collection began, and these are also included as deviations in the table.

### Subjects and subject preparation

Four subjects participated in the fMRI and pupillometry studies. All four participants are scientific investigators and three are authors of this study (4 males, ages 27, 28, 32, 46). These four participants choose to identify themselves by their initials. An additional 20 subjects, naïve to the hypotheses of the study, participated in the perception experiment (9 men, 11 women, mean age 27, range 20-33); their data have been assigned anonymous study identification labels. All subjects were screened for normal color vision^1^ and corrected acuity of 20/40 or better as assessed by the Snellen chart at a 20 foot distance. All subjects were studied at the University of Pennsylvania. The research was approved by the University of Pennsylvania Institutional Review Board and conducted in accordance with the principles of the Declaration of Helsinki. All subjects gave informed written consent.

Prior to fMRI scanning or perceptual rating, each subject underwent pharmacological dilation of the right eye (1% tropicamide ophthalmic with 0.5% proparacaine as a local anesthetic agent).

### Visual stimuli

We used the method of silent substitution with a digital light synthesis engine (OneLight Spectra) to stimulate targeted photoreceptors. Our device produces stimulus spectra as mixtures of 56 independent, ~16 nm full-width half-max primaries under digital control, and can modulate between these spectra at 256 Hz. Details regarding the device, stimulus generation, and estimates of precision have been previously reported.^2–4^

Our estimates of photoreceptor spectral sensitivities were as previously described,^3^ following the CIE physiological cone fundamentals.^5^ They account for the size of the visual field (64°), subject age, and the pupil size, which we assumed to be 8 mm in diameter under pharmacologic dilation.

Separate background and modulation spectra were identified to maximize available contrast on melanopsin and the combined stimulation of the L, M, and S cones. First, “mid-background settings were selected so as to maximize available melanopsin (or LMS) Michelson contrast for modulations symmetric around this background. Then, a 66.66% modulation was found. The negative ‘arm’ of this modulation served as the experimental background, and the positive ‘arm’ of this modulation represented the maximal, 400% contrast pulse. An additional constraint sought to minimize the difference in calculated chromaticity of the backgrounds of the LMS and melanopsin stimuli (Figures 1 and S1). The background for the LMS, Mel, and Splatter modulations were all nominally rod-saturating (100-200 cd/m^2^; >3.3 log sc td). The modulations did not explicitly silence penumbral cones.^3^

We elected not to perform psychophysical nulling of our stimuli for two reasons. First, in an earlier study^2^ we found that the test-retest reliability of nulling values produced by individual observers was not high. We estimated that stimulus adjustment for individual subjects was more likely to worsen photoreceptor silencing than to improve it. Second, we found that allowing for stimulus adjustment would reduce the available gamut in our modulations, with the consequence of a substantial reduction in available contrast on melanopsin.

We measured the melanopsin 400% background and stimulation spectra for a reference observer (32 years) before and after each scanning session for each subject during our initial fMRI experiment (described as Experiment 1 below). We calculated the average post-receptoral contrast for each of these 8 spectra (4 subjects x 2 measurements) with respect to the cone fundamentals assumed for the reference observer. From these measurements, we derived 8 sets of post-receptoral contrast values for LMS, L-M, and S− [L+M]. We then took the sign preserved absolute maximum value across each of the sets of 8 measurements. The resulting post-receptoral contrast values [%] were LMS: +2.173; L-M: +0.877; S-[L+M]: −10.451. Converted to cone contrast values [%] these were L: +3.050; M: +1.296; S: −8.278. We term this set of cone contrasts the 1x splatter control modulation. When presented to individual observers, the splatter control modulation was tailored to the age of the individual observers, such that even though the spectra seen by the observers would be different, they would all see a modulation with the nominal contrast values above, calculated using their age-corrected cone fundamentals.

The stimulus was viewed within an MRI compatible eye piece that provided a circular, uniform field of 64° diameter. The central 5° diameter was obscured. Subjects were asked to maintain fixation in the center of this obscured region to avoid stimulation within the macula, where spatial variation in macular pigment could alter the spectral properties of the stimulus.

Three-second pulses of spectral change were presented during individual trials of 16 s duration. During each trial, a transition from the background to the stimulation spectrum would occur starting at either 0, 1, or 2 seconds after trial onset (randomized uniformly across trials); this jitter was designed to reduce the ability of the subject to anticipate the moment of stimulus onset and to render trial timing asynchronous with respect to BOLD fMRI image acquisition. The transition from the background to the modulation spectrum, and the return to background, was subjected to a 500 ms half-cosine window.

The half-cosine windowing of the stimulus was designed to minimize perception of a Purkinje tree percept in our uniform spatial stimuli.^3^ Consider that there are both penumbral cones (that receive the stimulus spectrum after filtering through retinal blood vessels) and open-field cones, that receive the un-filtered stimulus. We have found previously that we can induce a percept of the retinal blood vessels using a uniform-field stimulus when two conditions are met: First, there is spatial contrast between the penumbral and open-field cones, and second, this spatial contrast is modulated at 4 Hz and higher.^3^ Both the LMS and melanopsin-directed spectral stimuli produce differential spatial contrast on the penumbral and open-field cones (on the order of 2-5%), satisfying the first condition. Critically, however, the Purkinje tree percept is ameliorated for these stimuli when modulated at 4 Hz and below. We windowed our stimuli with a 500 msec half-cosine at onset and offset. This corresponds to a 1 Hz modulation, and is thus comfortably below the slew rate that we have observed is needed to produce a spatial Purkinje percept.

### Simulation of biological variability causing inadvertent cone contrast

To address the amount of inadvertent stimulation of the L, M and S cones due to biological variability not captured by the CIE model for the cone fundamentals (Figure S5), we performed simulations of colorimetric observers, assuming variability in the following eight parameters: lens density, macular pigment density; L cone photopigment density, M cone photopigment density, S cone photopigment density; and the peak absorbance λ_*max*_ of the L, M and S cone photopigments. Using previously published estimates of the standard deviations in those parameters^6^, we randomly sampled independently from normal distributions with those SDs. The SDs were ±18.7% deviation in lens density, ±25% in macular pigment density; ±9% deviation in L cone density, ±9% deviation in M cone density, ±7.4% deviation in S cone density; and ±2 nm, ±1.5 nm and ±1.3 nm in λ_*max*_ for L, M and S cones respectively. Note that the variation in lens density was taken around the age-appropriate mean density for each subject. We performed this resampling 1,000 times, generating 1,000 sets of spectral sensitivities. This was done for the four observers from the fMRI studies (Figure S5) and the twenty observers from the perceptual studies (Figure S11).

We present plots of the L, M, and S cone contrasts after transformation to a post-receptoral opponent representation assuming mechanism sensitivities to cone contrast for luminance, red-green, and blue-yellow mechanisms of [0.5 0.5 0], [0.5 −0.5 0], and [−0.5 −0.5 1] respectively. This transformation corresponds to the DKL opponent color space representation^7^ when the background produces equal excitations in the L, M and S cones, for the case in which the L and M cone spectral sensitivities are scaled so that they sum to produce the luminous efficiency curve. We regarded this as a a reasonable choice of reference conditions to define the transformation, as it leads to intuitively straightforward properties of the assumed post-receptoral mechanisms. We note that for other backgrounds, this transformation will describe the opponent mechanism responses to the extent that those responses are the same for modulations seen against different backgrounds, when the LMS cone contrasts of the modulations are matched across backgrounds.

### Design of MRI experiments

Each of the four primary subjects participated in six MRI experiments (except for subject ASO who was unavailable to participate in the final, sixth experiment). The first two experiments presented pulses of either 400% melanopsin contrast only (Experiment 1) or 400% LMS contrast only (Experiment 2). Three experiments presented intermixed trials of different intensity of either “splatter” cone contrast (Experiment 3), melanopsin contrast (Experiment 4), or LMS contrast (Experiment 5). The final experiment (described in Figure S6) presented blocks of flickering L−M and melanopsin / rod modulations under scotopic and photopic conditions. During Experiments 1, 2, 3, and 6, the left eye was covered with an opaque patch. During Experiments 4 and 5 the left eye was uncovered and monitored with an infrared video camera (described below).

Each experimental session was approximately two hours. A given scanning session examined a single stimulus type (e.g., LMS, melanopsin, or splatter). The subject maintained adaptation to the background spectrum between trials and scan runs. Prior to each fMRI session, subjects underwent monocular dark adaptation for 20 minutes by wearing swimming goggles with the right eye obscured. Once in the scanner, the right eye was adapted for at least five minutes to the stimulus background prior to the start of functional scanning.

Experiments 3, 4 and 5 sought to measure the contrast response function (CRF) that related stimulus contrast to BOLD fMRI response. A set of stimuli of varying contrasts were presented in an intermixed order during a given scan. The LMS and melanopsin CRF studies presented 5, logarithmically spaced contrast levels (25, 50, 100, 200, and 400% contrast); the splatter CRF study presented 4 levels 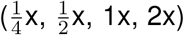. The 2x stimulus was in fact 1.95x due to limitations in device gamut; we adopt the technically inaccurate label for ease of description and interpretation. Ordering of these trial types within and across scans followed a pseudo-random, counter-balanced order.^8^

Functional MRI data collection took place during individual scans of 336 s duration. Between 9 and 12 scan runs were collected for each subject for each experiment. With the exception of Experiment 6, each scan run presented 21, 16 s trials; Experiment 6 presented blocks of stimulation and is described in Figure S6. Eighteen of the trials presented a spectral pulse. Three randomly selected trials presented an “attention event” instead of a stimulus pulse, during which the stimulus field dimmed for 500 ms. The subject was asked to press a button on a response pad when these dimming events occurred. Subjects performed well on this detection task. Collapsing performance across subjects and experiments, there were 0 false alarm responses during the 3,816 stimulus trials, and 11 misses during the 636 attention trials.

### MRI data acquisition and initial processing

MRI scanning parameters made use of the Human Connectome Project LifeSpan protocol (VD13D) implemented on a 3-Tesla Siemens Prisma with a 64-channel Siemens head coil. A T1-weighted, 3D, magnetization-prepared rapid gradient-echo (MPRAGE) image was acquired for each subject in axial orientation with 0.8 mm isotropic voxels, repetition time (TR)=2.4 s, echo time (TE)=2.22 ms, inversion time (TI)=1000 ms, field of view (FoV)=256 mm, flip angle=8°. BOLD fMRI data were obtained over 72 axial slices with 2 mm isotropic voxels with multi-band=8, TR=800 ms, TE=37 ms, FOV=208 mm, flip angle=52°. Head motion was minimized with foam padding. Although continuous pulse-oximetry was recorded, this physiologic measurement was not used in the fMRI data analysis.

The FreeSurfer (v5.3) toolkit (http://surfer.nmr.mgh.harvard.edu/)^9–12^ was used to process anatomical MPRAGE images to construct inflated brain surfaces and register data from across subjects for surface visualization. Briefly, this processing includes spatial inhomogeneity correction, non-linear noise-reduction, skull-stripping,^13^ subcortical segmentation,^14,15^ intensity normalization,^16^ surface generation,^9,10,17^topology correction,^18,19^surface inflation,^10^ and registration to a spherical atlas.^11^

Raw echo-planar volumetric data were motion corrected using the FMRIB Software Library (FSL) toolkit (http://fsl.fmrib.ox.ac.uk/fsl/). Motion corrected functional volumes were co-registered to subject-specific anatomy in Freesurfer using FSL-FLIRT with 6 degrees-of-freedom under a FreeSurfer wrapper (`bbregister`).

### BOLD fMRI time-series analysis

The pipeline for the analysis of the BOLD fMRI time series is available on GitHub (https://github.com/gkaguirrelab/MRklar/releases/tag/v1.0.0). Noise regressors were derived from the left, right, third, and fourth ventricles, as well as white matter, brain stem white matter, and non-brain tissue. Binary masks of these regions were initially identified in a Freesurfer anatomical segmentation volume (`aseg.mgz`). After co-registering to the functional volume, these regions were eroded by two voxels (for the white matter mask) or a single voxel (for all other regions) to avoid partial volume contamination from grey matter. The first five principal components of the time-series data across all voxels in these regions were then used as regressors. The signal from white matter local to each voxel was obtained and regressed. To obtain the local white matter signal for each voxel, the mean time series of all white matter voxels within a 15 mm radius sphere was regressed from the time series of the voxel found at the center of the sphere. This local white matter procedure was modeled after the ANATICOR pipeline in AFNI.^20^ Twenty-four motion regressors were derived from the initial six parameters that result from motion correction.^21^ The effects of these nuisance covariates were removed from the time-series data by regression. Finally, the time-series was subjected to a high-pass Butterworth filter with a cut-off of 0.01 Hz.

The primary analyses of the study were conducted within a V1 region of interest. A cortical surface atlas^22^ was used to define a patch of V1 cortex corresponding to the radial eccentricity range of 5–25°. For each subject, the average, post-processed signal within this region (and across the two hemispheres) was obtained for each scan run in each experiment. The regional time-series data were analyzed within a non-linear temporal fitting engine (https://github.com/gkaguirrelab/temporalFittingEngine).

As the timing of stimulus events (both spectral pulses and attention events) were asynchronous with respect to image acquisition (TRs), we derived the average evoked BOLD fMRI response for each stimulus type using a Fourier basis set approach.^23^ This approach provided an accurate estimate of the underlying response not available from a simple averaging of the time-series data itself across trials. The 16 s following the onset of each event was modeled with 8 harmonic pairs (a sine and cosine), ranging in frequency from 0.0625 to 1 Hz. The fit of the Fourier basis set to the evoked response was then averaged across scan runs. Because of jitter introduced into the timing of onset of each event, the inter-stimulus-interval (ISI) ranged between 14 and 18 seconds. Because the stimulus order was counterbalanced, the additive effects of trial overlap (for when the ISI was <16 seconds) should be estimated efficiently and without bias by the model. Any non-linear interactions caused by hemodynamic response overlap will not be captured in our model, but we are reassured that there is nothing unusual in the appearance of the evoked response estimates between 14 and 16 seconds.

The evoked responses obtained in this way to the spectral pulse stimuli are presented in Figures 2d, 2e and S2. As the attention events were brief (500 msecs), the average evoked response to the attention events was taken as an estimate of the hemodynamic response function (HRF) for each subject (Figure S3a). We observed that our subjects differed in the overall amplitude of their BOLD fMRI HRF. We obtained the peak amplitude of the HRF for each subject, and then divided each value by the mean of the values across subjects. We treated the result as a “subject scaler” that was used to normalize subsequent measurements of response amplitude from each subject to remove this individual difference.

The evoked response for each spectral-pulse stimulus type for each subject was then modeled with a two-parameter model (Figure S3b). The first parameter controlled the duration of a step-function of neural activity that was then convolved by the HRF for the subject. The resulting shape of BOLD fMRI response was normalized to have unit amplitude, and then subjected to a gain parameter. The best fitting parameters (in the least-squares sense) were found by non-linear search (`fmincon`).

We conducted whole brain (cortical surface) analysis of the data from Experiments 1 (400% melanopsin only) and 2 (400% LMS only). The time-series data from each voxel for each subject was projected to hemisphere-symmetric cortical surface atlas (fsaverage-sym) and smoothed on the cortical surface using a 5 mm full-width at half-maximum Gaussian kernel. An approximation to a Fourier basis set analysis was conducted on the time-series data at each cortical point for each of the *k* scan runs using the FSL FEAT and FOBS routines, modeling the 14 s period following each stimulus event with a set of 14 sinusoids that varied in frequency from half a cycle per period to 14 cycles per period. The *p*-value associated with the F-statistic for this model was obtained at each vertex. For each subject and hemisphere (at each cortical point), the set of *p*-values across the *k* scan runs were used to calculate a *χ*^2^-value with 2*k* degrees of freedom using Fisher’s method. The map of *p*-values corresponding to the *χ*^2^map from each subject and hemisphere were combined again using Fisher’s method, and the resulting maps of *χ*^2^-values were used to illustrate the evoked stimulus effect shown in Figures 2a, b. These maps were thresholded at a value of *χ*^2^ (16 df)=61.4. This corresponds to a Bonferroni corrected, map-wise *p* = 0.05 threshold after accounting for the number of vertices in the group map (and disregarding map spatial smoothness).

### Eye and pupil tracking

Infra-red (IR) video eye-tracking was performed during Experiments 4 and 5. The LiveTrack AV MR-compatible eye tracking camera (Cambridge Research Systems, Rochester, UK) was used to record video from the left eye of each subject at either 60 Hz or 30 Hz (the lower frame rate was used in the Mel CRF studies for ASO and GKA, and in the LMS CRF study for ASO). The camera was attached to the 64-channel head coil using a custom mount, and positioned 10-15 cm away from the left eye of the subject. The camera and head coil were draped in black felt to minimize scattering of light to the left eye from the eyepiece over the right eye. Consistent with the minimization of scattered light and room light, subjects generally reported binocular suppression of the left eye during these experiments.

A live video feed from the system was used to monitor subject alertness and head motion during scanning. The system recorded the position of the IR glint on the tear film and the size and position of an ellipse fit to the outline of the pupil in each frame, and from this derived pupil size and eye position. We found that the automated ellipse fitting was unstable in the vertical dimension. For this reason, we report here the pupil size derived from the horizontal width of the fitted ellipse, and eye position in the horizontal plane only. The timing of data collection was synced with MRI scan acquisition and stimulus presentation using an analog signal (TTL) sent by the scanner at the start of the scan and at the time of each image repetition (TR). Absolute pupil size was determined by calibrating the camera against targets of known dimension following each scan session.

The analysis pipeline for the pupil and eye position data is available on GitHub (https://github.com/gkaguirrelab/pupilMelanopsinMRIAnalysis). First, blinks were identified as timepoints during which the glint was not visible. The pupil size and position measurements were set to NaN in the 50 ms before and after each blink. The pupil size vectors were then subjected to a 0.025 Hz high-pass filter. A 13 s period of pupil response following the onset of every trial was extracted, expressed in percent change units, and set to have a value of zero during the first 100 ms following the onset of the stimulus. The median response across trials for each stimulus type for each subject was obtained (Figure S8). Each median response was then fit with a six-parameter, three-component pupil temporal model (Figure S9) using a non-linear search (fmincon) within a temporal fitting engine (https://github.com/gkaguirrelab/temporalFittingEngine).

We observed that our subjects differed in the overall amplitude of their pupil response. For each subject, we obtained the total area of pupil response (% change x seconds) across all stimulus types (mel and LMS pulses of every contrast level). We divided each value by the mean of the set of values across subjects. We treated the result as a “subject scaler” that was used to normalize measurements of response amplitude from each subject to remove this individual difference.

### Perceptual rating experiment

Perceptual ratings were obtained from experimentally naïve subjects using the same stimulus presentation apparatus as was used in the fMRI experiments. Subjects were positioned in a chin rest in a darkened room and observed stimuli with their pharmacologically dilated right eye. The experiment was composed of several periods. In each period, the subject would first adapt to a stimulus background, and then view spectral pulses of a particular stimulus type (light flux, LMS, or melanopsin). Three initial “exposure” periods were used to familiarize subjects with the procedure and the perceptual range of the stimuli. Each exposure consisted of 1 min of adaptation to the background, followed by presentation of 3 spectral pulse trials of a given type. Subjects were asked only to observe the stimuli. Following the exposure periods, the subject participated in six “rating” periods. Each rating period consisted of a 5 min adaptation to a background, followed by the presentation of 9 spectral pulse trials of a given type. Before each trial, the subject was read a description of a perceptual property that they were to rate for the upcoming stimulus trial. Following presentation of the stimulus pulse, the subject was prompted for their rating on a scale of 1 to 7. The subject could ask for the description and pulse to be repeated one additional time prior to providing a rating. The subject was asked to rate a different perceptual property for each of the 9 trials in a given rating period. Rating periods for light flux, LMS, and melanopsin stimuli were each conducted twice, with subjects randomized to follow one of two trial orders:

i. light flux, melanopsin, LMS, light flux, melanopsin, LMS
ii. light flux, LMS, melanopsin, light flux, LMS, melanopsin

The nine perceptual properties were defined by pairs of antonyms (e.g., cool-warm) that defined the extreme ratings of 1 and 7. Subjects were instructed to fixate the center of the stimulus field and report the appearance of the light pulse in visual periphery, doing their best to ignore any percept within or adjacent to the obscured macular region.

The nine perceptual properties were defined by pairs of antonyms (e.g., cool-warm) that defined the extreme ratings of 1 and 7. Subjects were instructed to fixate the center of the stimulus field and report the appearance of the light pulse in visual periphery, doing their best to ignore any percept within or adjacent to the obscured macular region.

For the perceptual rating experiment, our photoreceptor spectral sensitivity estimates assumed a (27.5°) field for generating the receptor-isolating modulations while the observed field was in fact (64°) as in the fMRI experiments. This lead to numerical but insignificant differences in the estimate for the macular pigment density. In the contrast and splatter calculations for this experiment, we assumed the 64° in our estimates for the spectral sensitivities.

The first two dimensions of the principal components analysis of the perceptual rating data were used to describe the results as the addition of further dimensions was found to reduce cross-validated categorization accuracy.

### Spectrum seeking to improve stimulus control

The stimuli used in the perceptual study were subjected to an additional refinement prior to each data collection session, designed to further reduce inadvertent cone contrast in the melanopsin-directed stimulus. An adaptive spectrum-correcting procedure addressed uncertainty in our device calibration due to instrumental drift and small failures of primary additivity. This procedure adjusted the mirror settings in our digital light synthesis engine so as to match the nominal, receptor-isolating spectra. This procedure was performed for the age-adjusted stimuli of all subjects in the perceptual rating experiment.

We started with a pair of primary values designed to yield a certain contrast: The background primary values *P*_*BG*_ and the modulation primary values *P*_*Mod*_. The spectral calibration procedure of the light synthesis engine determines the primary matrix *M*, which, when multiplied with the primary values and added to the dark spectrum *spd*_*dark*_, yields the predicted target spectra *spd*_*BG; target*_ and *spd*_*Mod; target*_. Contrast properties of the stimulus are defined with respect to these two spectra.

During a validation, we gamma-correct the linear primaries values, *P* using our device calibration model. This yields the pair of settings *S*_*BG*_ and *S*_*Mod*_. These are then provided to the light engine and spectral measurements *spd*_*BG; val*_ and *spd*_*Mod; val*_ are obtained. Due to imprecision in the stimulus control, *spd*_*BG; target*_ and *spd*_*BG; val*_, and *spd*_*Mod; target*_ and *spd*_*Mod; val*_, are different.

The goal of the adaptive procedure is to find terms ∆*P*_*BG*_ and ∆*P*_*Mod*_ which correct the primary values. To do this, we do the following (*i* ϵ 1 … *N*, where typically *N* = 10).

i. Gamma correct: *P_i; BG_* → *S_i; BG_* and *P_i;_ Mod* → *S_i; Mod_*
ii. Obtain target spectra: *P_1_, _BG_* → *spd_BG; target_* and *P_1, Mod_* → *spd_Mod; target_*
iii. Measure *spd_i; BG; val_* and *spd_i; Mod; val_*
iv. Calculate the spectral difference between target and validated spectra in the *i*-th iteration: ∆*spd_i; BG_ = spd_BG; target_ − spd_i; BG; val_* and ∆*spd_i; Mod_ = spd_Mod; target_ − spd_i; Mod; val_*.
v. Obtain ∆*P_i; BG_* and ∆*P_i; Mod_* corresponding to the spectral differences using our calibration model that maps spectra back to device coordinates.
vi. Update the primary values for the next iteration: *P_i+1; BG_ = P_i; BG_ + λ∆P_i; BG_* and *P_i+1;_ Mod = P_i; Mod_ + λ∆P_i; Mod_*, where λ = 0.8 is the learning rate.

We find that this spectrum-correction procedure reliably reduces inadvertent cone stimulation due to uncertainty in device control.

### Data and code availability

All raw data are available as packaged and MD5-hashed archives as well as tables detailing the biological variability on FigShare (https://figshare.com/s/0baea6ed50758abbabf4). All code is available in public GitHub repositories (https://github.com/gkaguirrelab/Spitschan_2017_PNAS/). Un-thresholded statistical maps from Experiments 1 and 2 for each subject are available from NeuroVault (http://neurovault.org/collections/2459/).

## Supplementary Material

**Figure S1:**
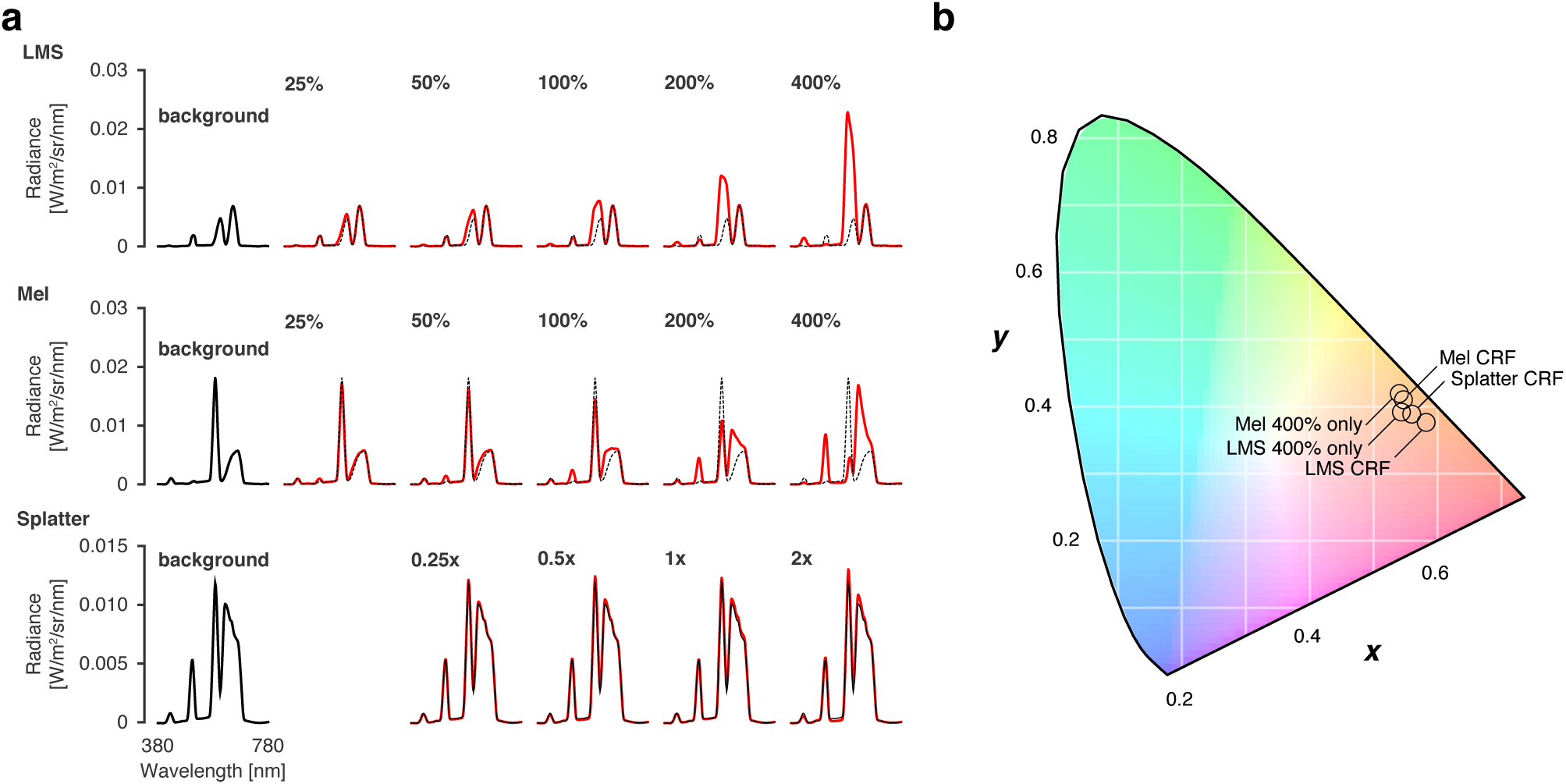
Spectra and chroma of all stimuli (related to Figure 1). **(a)** The stimulation spectra (red) for each contrast level in comparison to the background spectrum (black) for the LMS, melanopsin, and splatter stimuli. **(b)** The calculated CIE 1931 chromaticity^1^ for all stimulus backgrounds. Experiments that presented stimuli of different contrast levels are indicated with “CRF” (Contrast Response Function).

**Figure S2:**
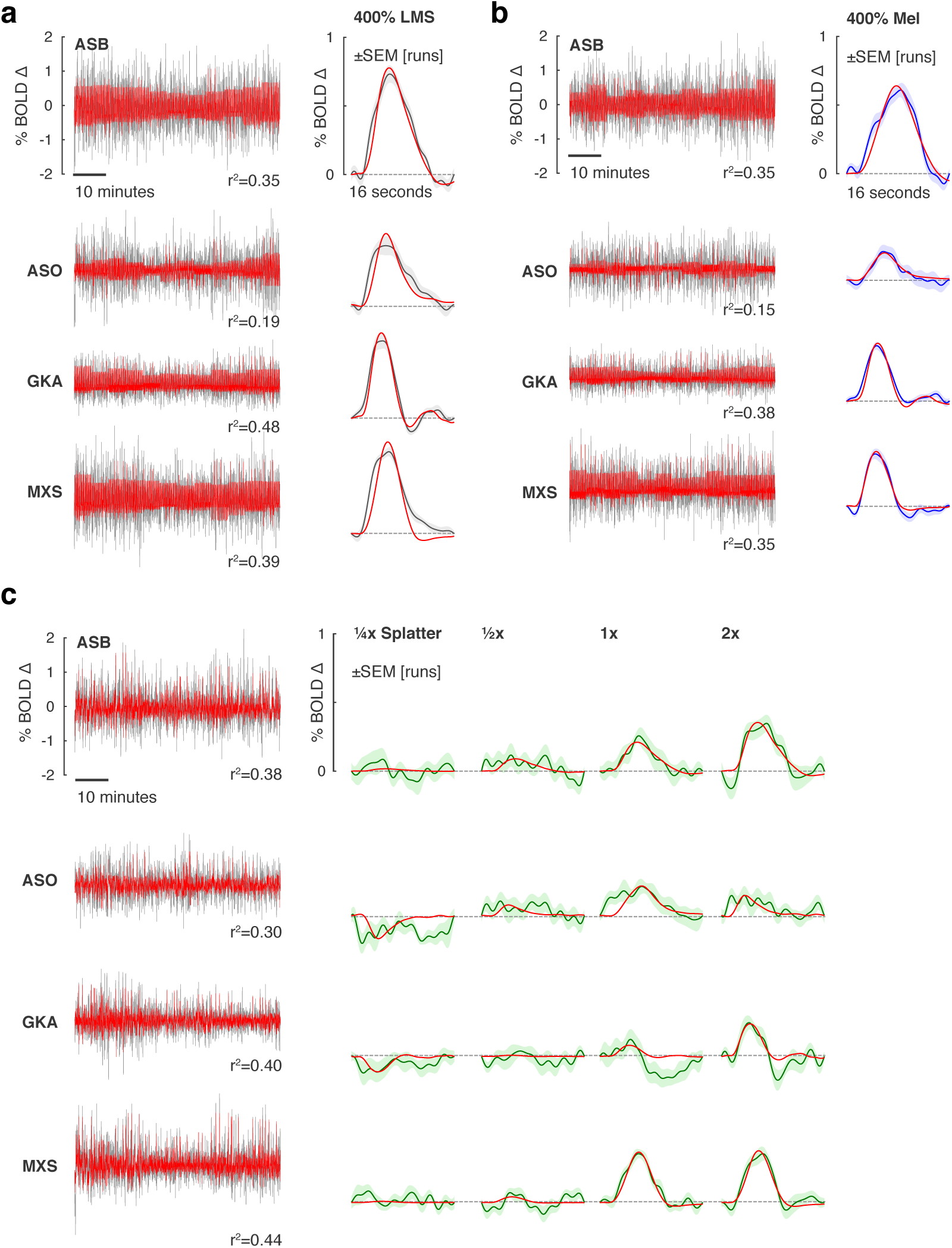
Additional BOLD fMRI time-series and model fits (related to Figure 2). **(a)** V1 responses to 400% LMS stimulation. Our initial study to explore broad cortical responses presented only trials with 400% stimulus contrast. *Left* The BOLD fMRI time-series data from the area V1 region for each subject (black), following preprocessing to remove nuisance effects. A Fourier basis set modeled (red) the mean evoked response to each contrast level during each run with the *r*^2^ values of the model fit indicated. *Right* The evoked responses for each subject to the 400% LMS stimuli (black), and SEM of the response across the 9-10 scanning runs performed in each subject (shaded region). The responses were fit by a model (red) that convolved a step function of neural activity by the hemodynamic response function measured for each subject. **(b)** The corresponding responses within the V1 region to melanopsin stimulation of 400% contrast. **(c)** The corresponding responses within the V1 region to the “splatter” modulation, with contrast varying from one-quarter to two-times the estimated cone splatter contrast arising from device imprecision (see Online Methods, Figure S5).

**Figure S3:**
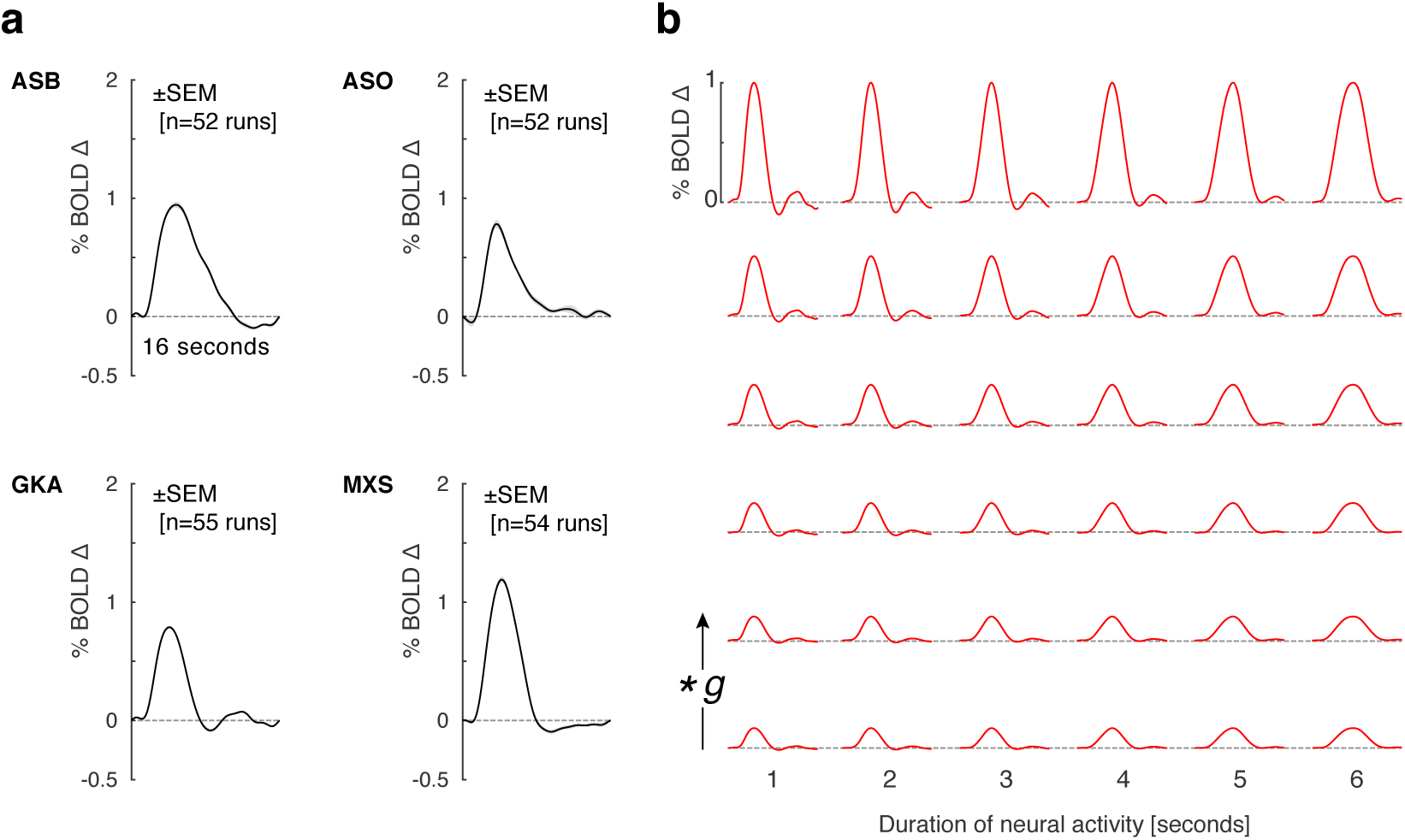
HRFs and evoked response model (related to Figure 2). **(a)** In all experiments, 14% of stimulus trials were randomly replaced with an attention event, during which the stimulus dimmed for 500 ms and in response to which the subject was to press a button on a response pad. The same response events occurred in each of the >50, 336 second scan runs for each subject across all experimental conditions. The BOLD fMRI response evoked within the studied V1 region in response to the attention events was estimated using a Fourier basis set for each run for each subject. The 16 s that followed each event was modeled with 8 harmonics, providing a temporal resolution of 1 Hz. The average response across runs (black) for each subject (expressed in units of percent BOLD signal change) was taken to be an estimate of the hemodynamic impulse response for that subject and was used in modeling of fMRI responses to other stimulation conditions for that subject. The SEM of the response across runs (shaded gray) is in most cases smaller than the plot line. **(b)** Shown are how the predicted BOLD fMRI responses for subject GKA vary with inferred duration of neural activity (across each row) and amplitude of BOLD fMRI response (each row shows a different value of *g*). The model varied the duration of a step function of neural activity that was then convolved with the HRF for that subject and subjected to multiplicative scaling (\**g*) to best fit the evoked response. The fits provided by this model are shown in Figure 2 and Figure S2, and the amplitude and duration parameters derived from fitting are the subject of Figures 3 and S4.

**Figure S4:**
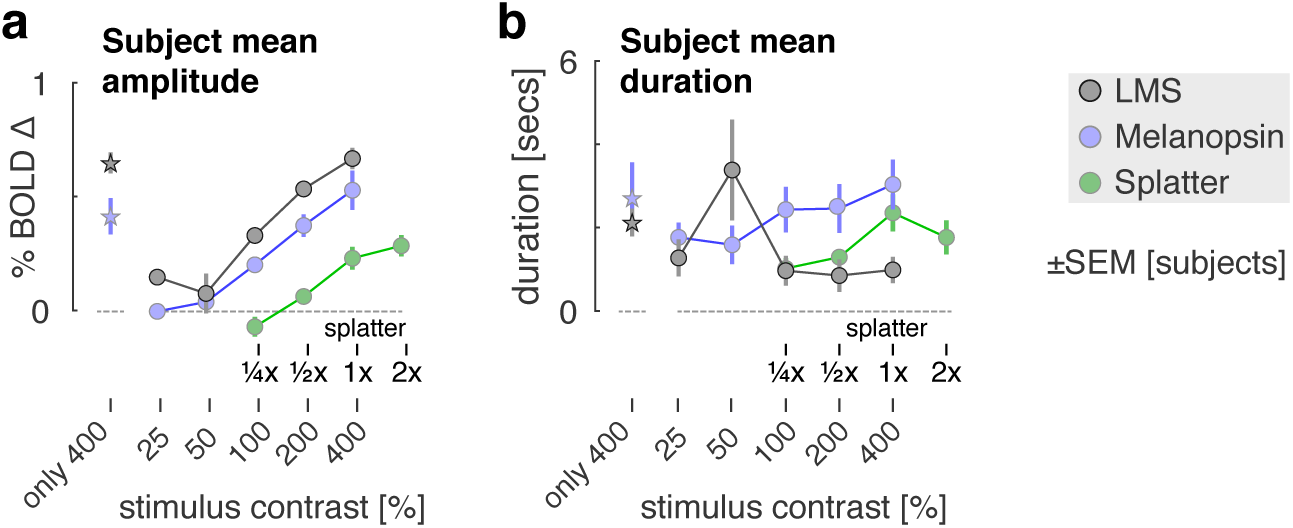
Amplitude and duration of response in V1 by stimulus contrast (related to Figure 3). **(a)** The mean amplitude of evoked response with the V1 region across subjects for each contrast level is shown for the LMS (gray), melanopsin (cyan), and “splatter” (green) stimulus conditions. The star symbols are the amplitude measurements obtained in the initial, 400% contrast only LMS and melanopsin studies. The 1x splatter condition presented cone contrast equal to the maximum inadvertent contrast measured in validated spectra in the melanopsin and LMS experiments. We calculated as well the amplitude of response for the 400% contrast only LMS and melanopsin studies within visual areas V2 and V3 (for which we have available an eccentricity map from cortical anatomy^2^). Within area V1, the response amplitudes (± across subjects) were 0.6459 ± 0.1000 and 0.4123 ± 0.1799 for LMS and melanopsin, respectively. Within V2 the values were 0.2675 ± 0.1177 and 0.3684 ± 0.0842 (LMS and melanopsin), and within V3 they were 0.2531 ± 0.1242 and 0.3790 ± 0.0657 (LMS and melanopsin). Overall, the response to wide-field LMS stimulation declined across visual areas (as reported previously^3^) while the response to melanopsin was more evenly maintained across these early visual areas. **(b)** The mean modeled duration across subjects of underlying neural activity within the V1 region is shown for the three stimulus conditions.

**Figure S5:**
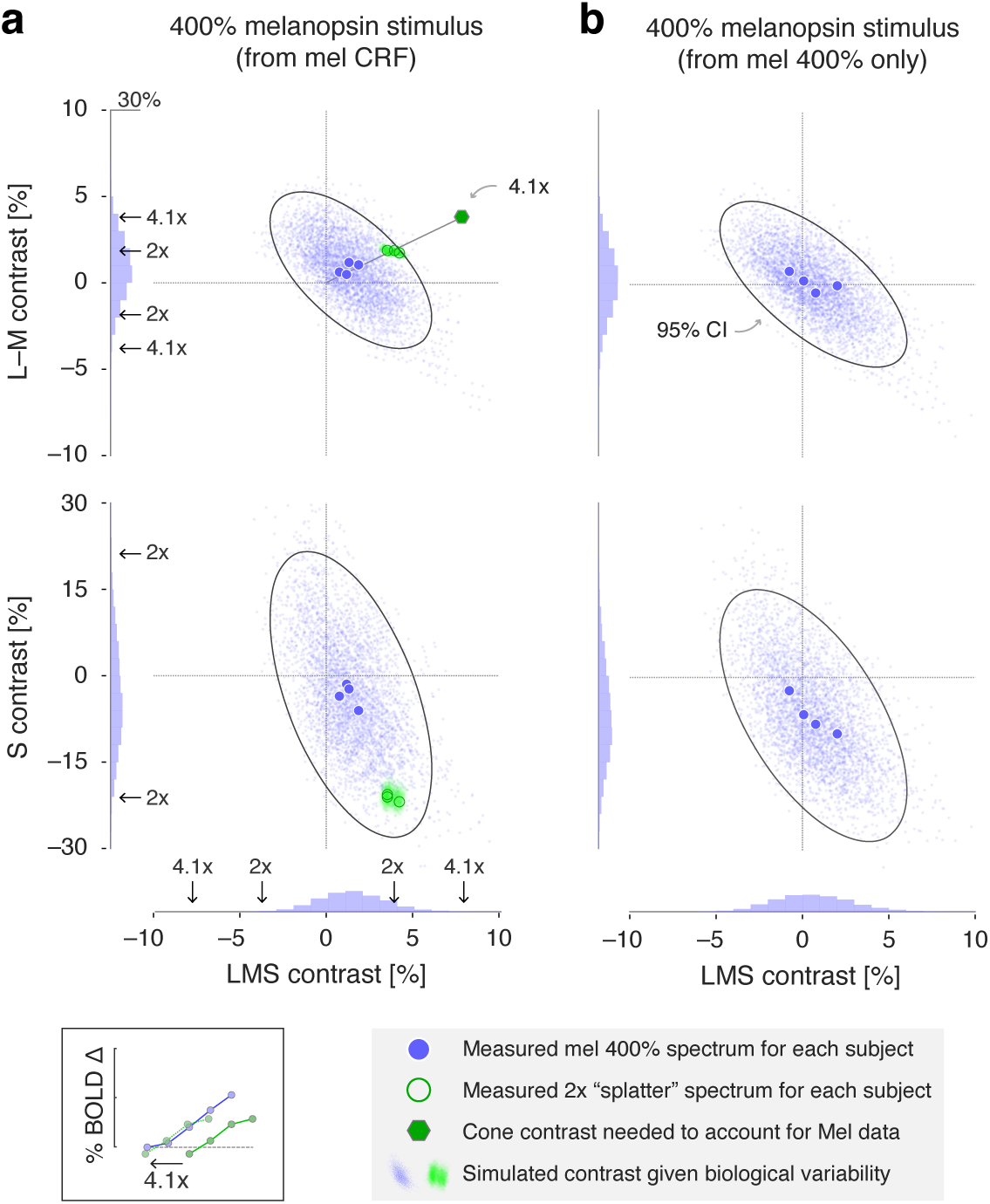
Inadvertent cone contrast in the fMRI stimuli (related to Figure 3). **(a)** Due to biological variability and inevitable imperfections in device control, a nominally cone silent modulation will produce inadvertent contrast upon the cones. We considered the extent to which this undesired contrast could account for the BOLD fMRI signals we observed in response to a melanopsin-directed spectral pulse. For each subject, multiple measurements of the 400% melanopsin-directed stimulus spectrum were made before and after each data collection session. This set of measurements was averaged for each subject to produce a single spectrum, which was then submitted to a calculation (https://github.com/spitschan/SilentSubstitutionToolbox) that estimated the degree of contrast upon each of the post-receptoral cone mechanisms (L−M, S, LMS). The four large, blue circles in each plot indicate the calculated contrast caused by device imprecision for the stimuli seen by each of the observers. We created a stimulus modulation (“1x splatter”; Figure S1a) that had cone contrast equal to the max, across-subject contrast attributable to device imprecision. A set of “splatter” stimuli with log-spaced intensity 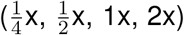 were derived from this initial modulation and studied during a control BOLD fMRI experiment. The spectrum of the 2x modulation was measured for each experimental session for each subject, and the cone contrast estimated in this modulation is indicated by the large, green circles (one circle for each observer; some plot symbols are overlapped). We next considered how biological variability could cause these estimates of cone contrast to change. Our model of cone contrast incorporates assumptions regarding: lens transmittance; density of macular pigment; L, M, and S cone density; and variation in the peak spectral sensitivity (λ_*max*_) of the L, M, and S cones. We simulated biological variation in these parameters by conducting 1,000 re-calculations of the cone contrast for each subject, using values for each parameter drawn from published distributions of individual differences.^4^ The cone contrast returned by each simulation comprises a point in the cloud of blue values in each plot; an ellipse (solid line) indicates the iso-probability contour that encloses 95% of the 2D projection of the boostrapped values upon the post-receptoral axes, computed assuming that the underlying distribution was a bivariate Gaussian. The marginal distribution of this set of simulated contrast values is shown on each cardinal axis. The same calculation was conducted for the 2x splatter spectra, yielding the cloud of green points. We next related these values to our BOLD fMRI measurements. We have for each subject a contrast response function (CRF) for melansopin and for multiples of inadvertent cone contrast (splatter) due to device imprecision (Figure 3). For each subject, we asked how much larger the splatter contrast would have to have been to produce responses that match the melanopsin CRF; this amounts to asking how many log-units the splatter CRF must be shifted to the left to best match the melanopsin CRF (inset, bottom left). Across subjects, the mean shift multiplier was 4.1 (individual values were ASB 3.2, ASO 2.8, GKA 7.7, MXS 4.1). Extending the line that connects the origin of the cone-contrast space and the 2x splatter modulation, we identified the position that would correspond to a 4.1x splatter modulation (green hexagon). We considered the position of this point (and its mirror symmetric reflections) in the opponent modulation space with respect to the marginal distributions of simulated inadvertent contrast due to biological variability and device imprecision. The key observation is that the inadvertent cone contrast necessary to produce the observed BOLD fMRI responses to the 400% melanopsin stimulus are unlikely to have occurred. The proportion of simulated contrast values (in both tails) that exceed the 4.1x level is 0.2% on the LMS dimension; 0% on the S dimension; and 5.3% on the L−M dimension. To account for our data, one or more of these values would have to have been exceeded for all four subjects. The odds of this occurring for a single subject is: *P*(LMS or L-M exceeded) = 1 − ((1 − 0.053) × (1 − 0.002)) = 0.0549 and the odds of this occurring for all four subjects is *p* = 9.1 ×10^−6^. **(b)** The corresponding calculation of cone contrast due to device imprecision and biological variability for the melanopsin stimulus used in the 400% contrast only experiment.

**Figure S6:**
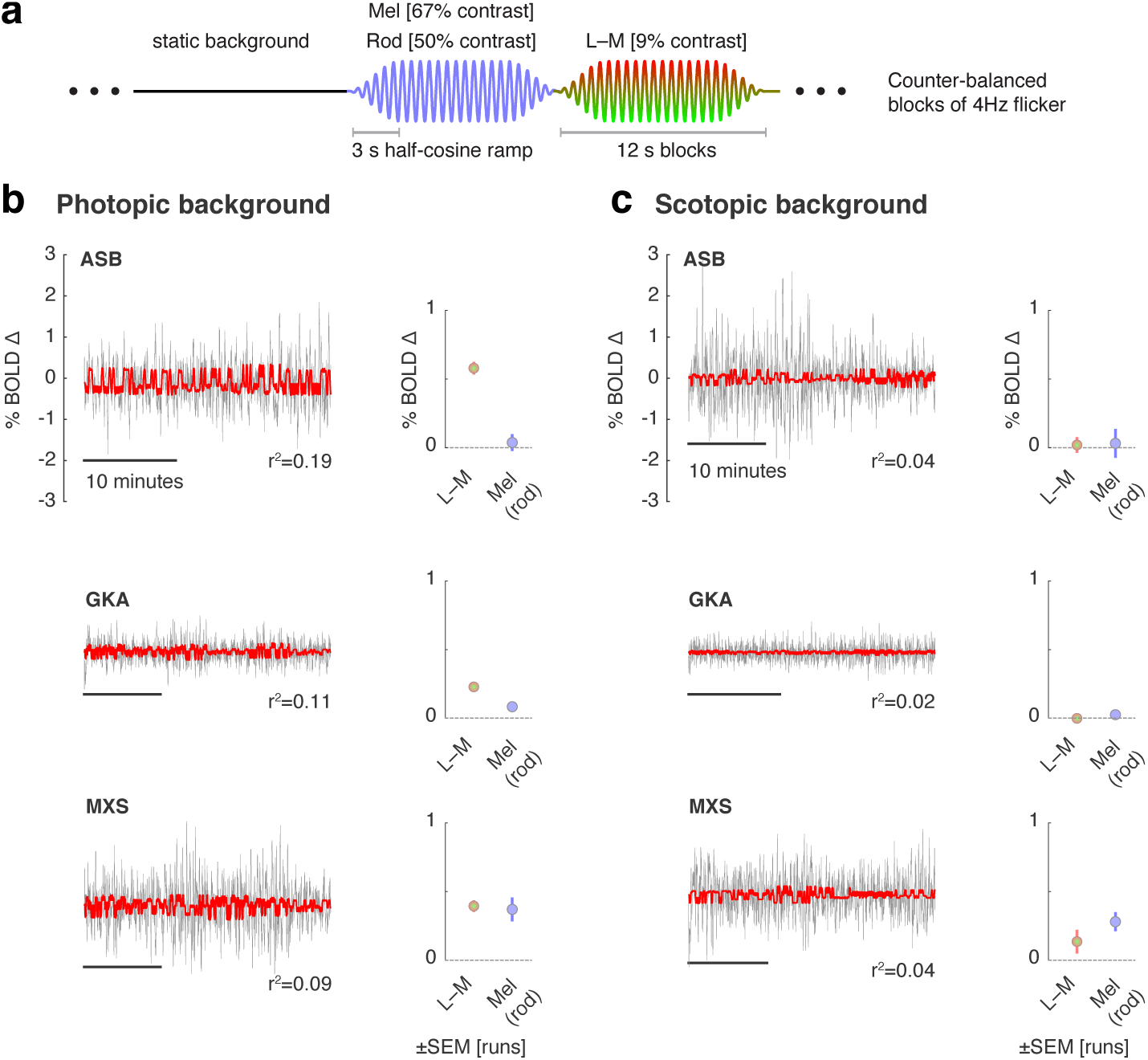
An unsuccessful control experiment (related to Figure 3). The rod and melanopsin spectral sensitivity functions overlap extensively. The background used for our melanopsin directed stimuli was 3.5 log_10_ scotopic Trolands (scot Td), nominally at or above the rod saturation threshold, found to be 3.0 log_10_ scot Td (Figure 2 of Adelson 1982)^5^ or 3.3-3.7 log_10_ scot Td (Aguilar & Stiles 1954).^6^ Therefore, we may expect in our experiments that there is no, or minimal, time-varying signal contributed by the rods. Nonetheless, we considered control experiments that could address the possibility of rod intrusion. While it is possible in principle to create a melanopsin directed stimulus that silences both the rods and cones, in practice we find that our device is limited to a maximum 60% unipolar (Weber) contrast pulse directed at melanopsin while silencing both rods and cones. Given our finding that at least 100% unipolar melanopsin contrast is needed to produce a reliable cortical response, we regarded this stimulus as ineffective. Instead, we examined whether the response to our melanopsin directed stimulus varied as a function of temporal frequency, with the logic that melanopsin responses would be attenuated to a stimulus modulated at 4 Hz, while rod responses would persist. Ultimately we found this experiment to be uninformative. The BOLD responses evoked by the stimuli were small and / or poorly modeled, with low r^2^ values, particularly in the scotopic condition. Moreover, inconsistent responses were obtained across subjects. Despite our inability to draw clear conclusions from these measurements, we present the data here for completeness. **(a)** The experimental design was adapted from a prior study.^7^ Around a common background, we presented a 4 Hz modulation that targeted either L−M with a 9% bipolar (Michelson) contrast (while silencing the rods) or melanopsin with 67% bipolar contrast on melanopsin and 50% bipolar contrast on rods. The modulations were presented in 12 s blocks, with a 3 s half-cosine window at onset and offset, in a counter-balanced order. **(b)** Photopic conditions. *Left* The BOLD fMRI time-series data from the area V1 region for each subject (black), following pre-processing to remove nuisance effects. The data were modeled (red) with a step-function for each stimulus condition, convolved by subject-specific hemodynamic response function. *Right* The amplitude of evoked responses for each subject for the 4 Hz L−M and melanopsin modulation blocks as compared to the static background. **(c)** The corresponding data obtained during scanning under scotopic conditions. Subjects dark-adapted for at least 20 minutes prior to scanning. A 6 log unit neutral density filter was placed in the light path, reducing the stimulus background to approx. 0.0001 cd/m^2^.

**Figure S7:**
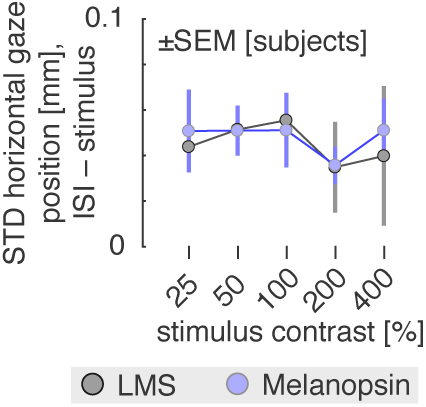
Variation in horizontal gaze position with stimulation (related to Figure 4). Subjects were asked to maintain fixation upon the center of a 5° opaque circle. Infrared video of the left eye was recorded during functional MRI scanning in some experiments. We measured the horizontal position of the eye during the scanning session to examine if stimulus presentation led to systematic changes in fixation stability. While vertical eye position was recorded, these data were not considered given that the eye has less fixational variation in the vertical plane, and the generally noisier quality of the vertical position data. The standard deviation of eye position was measured during the three seconds of stimulus presentation and during the ensuing interstimulus interval (ISI). The mean difference (averaged across subjects) between the ISI and stimulated periods was obtained for the LMS (gray) and melanopsin (blue) stimuli at each contrast level. A clear effect of stimulation was to reduce horizontal eye movement as compared to the ISI period (all data points different from zero). This effect did not systematically vary by stimulus type (LMS or melanopsin) or by contrast. Therefore, differences in BOLD fMRI responses between contrast levels or stimulus type are not explained by differences in evoked eye movements. It remains possible, however, that measured cortical responses to stimulation contain some constant component of change in eye movements. This effect may contribute to prior results.^8^

**Figure S8:**
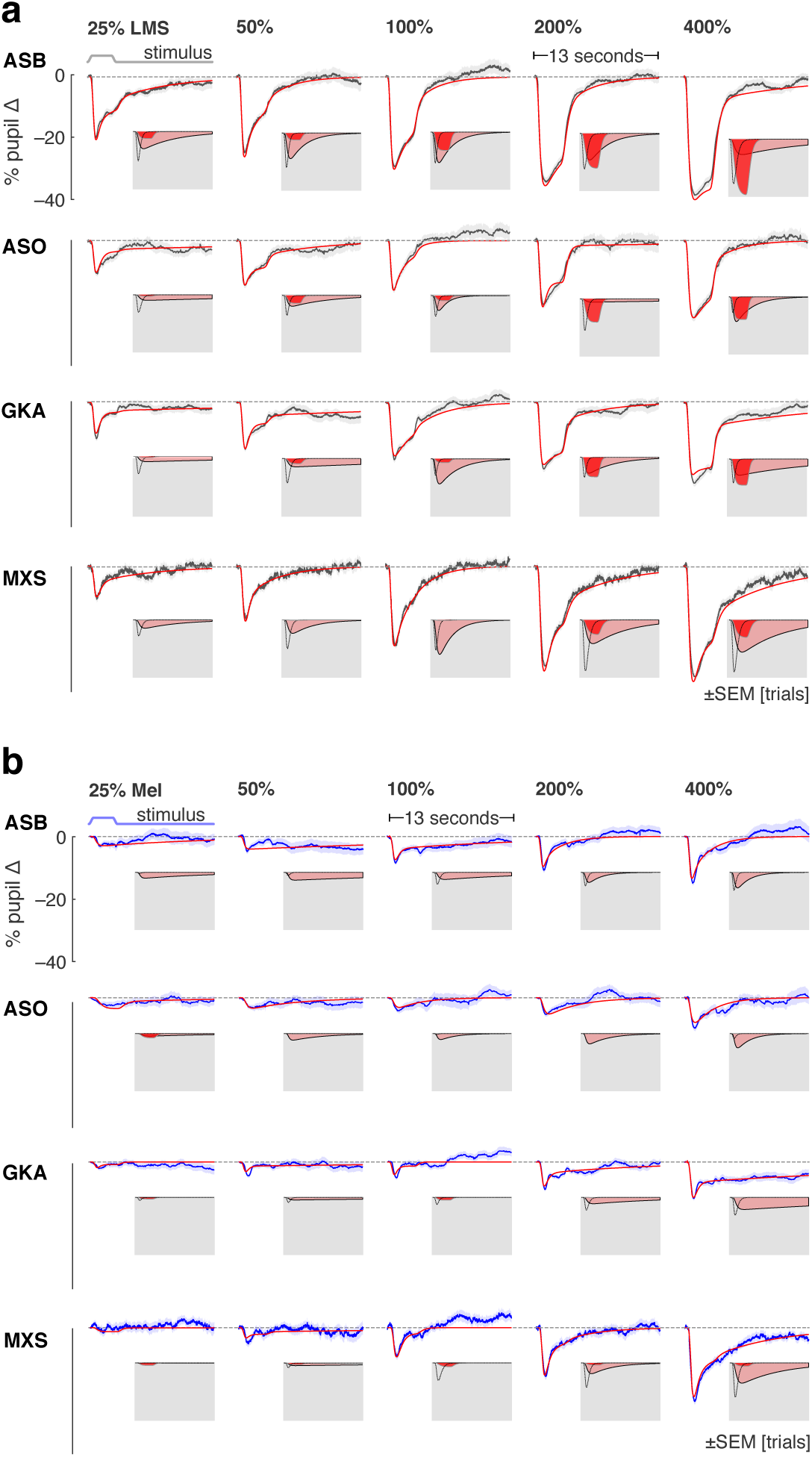
Individual subject pupil responses (related to Figure 4). The consensual pupil response of the left eye was measured during stimulation of the pharmacologically dilated right eye. **(a)** The mean (across trials) pupil response evoked by LMS stimulation of varying contrast levels (black), with SEM across trials (shaded). Each row contains the data from a different participant. The evoked response was fit with a three component, six-parameter model (red). The three components that model each response are shown inset on a gray field. **(b)** The corresponding mean pupil responses evoked by melanopsin stimulation of varying contrast levels.

**Figure S9:**
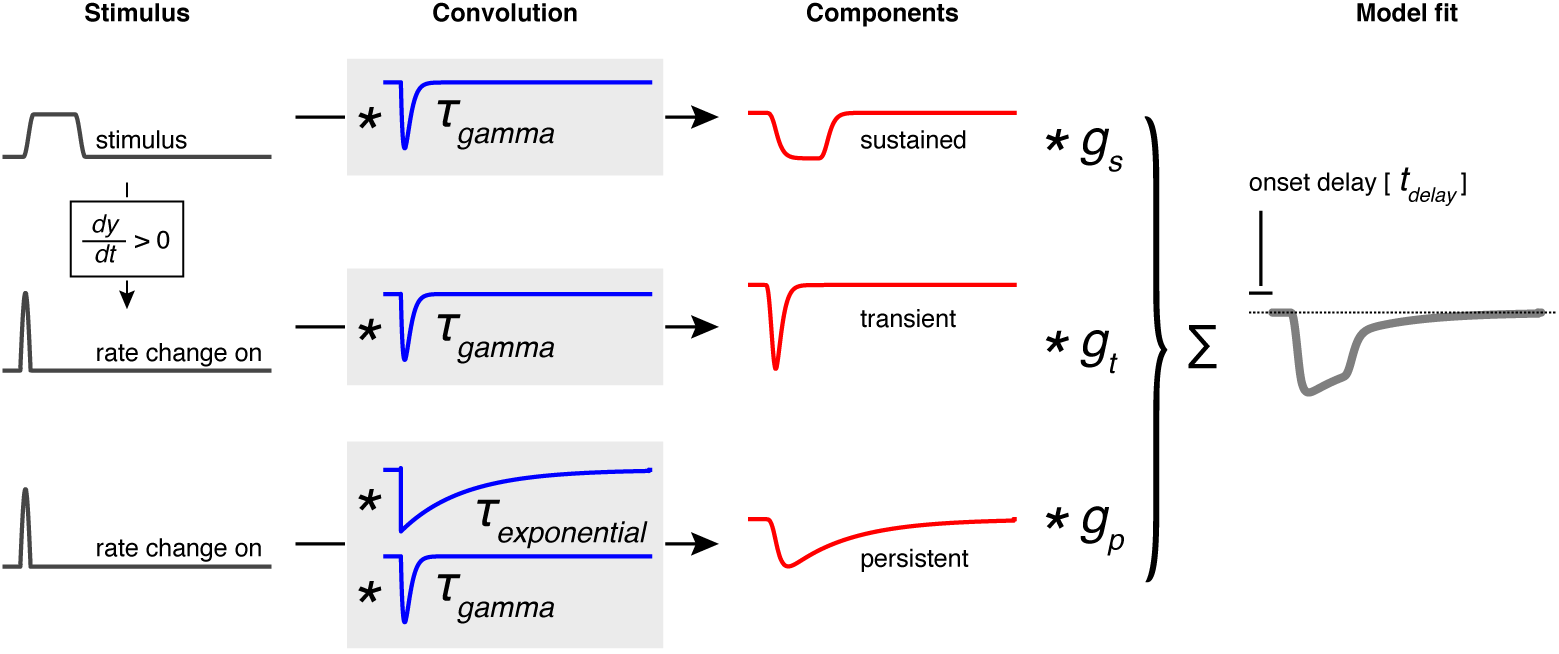
Pupil temporal model (related to Figure 4). The across-trial, within-subject average evoked pupil response to each stimulus type (LMS and melanopsin) and contrast level was fit with a six-parameter, three-component model using a non-linear temporal fitting engine (https://github.com/gkaguirrelab/temporalFittingEngine). The model was designed to capture the three, visually apparent and temporally separated components of the evoked pupil response. The elements of the model are not intended to directly correspond to any particular biological mechanism. The input to the model was the stimulus profile (black). An additional input vector, representing the rate of stimulus change at onset, was created by differentiating the stimulus profile and retaining the positive elements. These three vectors were then subjected to convolution operations composed of a gamma and exponential decay function (blue), each under the control of a single time-constant parameter (*τ_gamma_* and *τ_exponentiai_*). The resulting three components (red) were normalized to have unit area, and then subjected to multiplicative scaling by a gain parameter applied to each component (*g_transient_*, *g_sustained_*, and *g_persistent_*). The scaled components were summed to produce the modeled response (gray), which was temporally shifted (*t_delay_*). We observed that some evoked responses for some subjects had a late dilation phase in which the pupil became larger than its baseline size. We did not attempt to capture this inconsistent behavior in our model.

**Figure S10:**
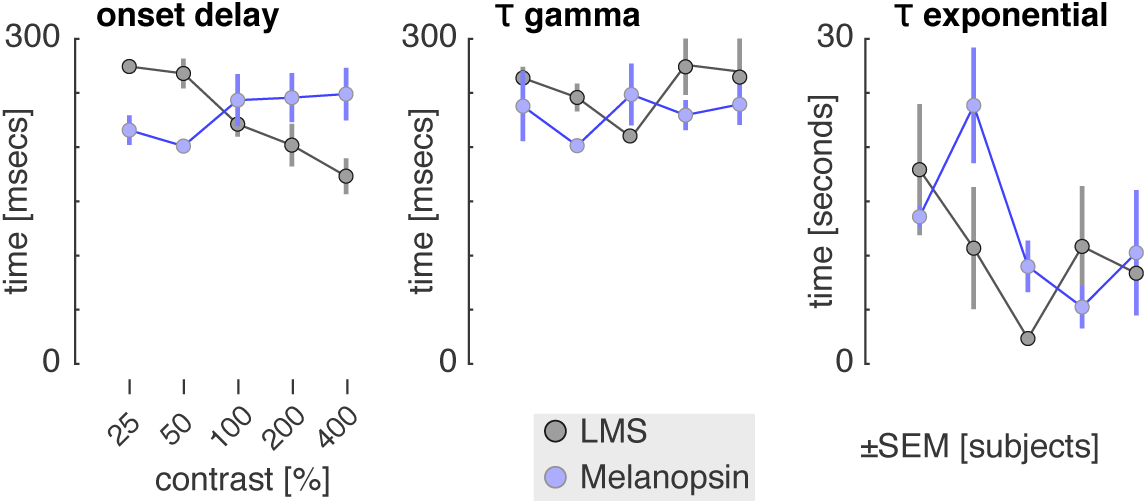
Temporal pupil model parameters by contrast (related to Figure 4). Pupil responses were fit with a six-parameter model, of which three parameters controlled the temporal behavior of the model. Each plot presents the mean (across subjects) of a temporal parameter, as a function of contrast for LMS (gray) and melanopsin (blue) stimulation.

**Figure S11:**
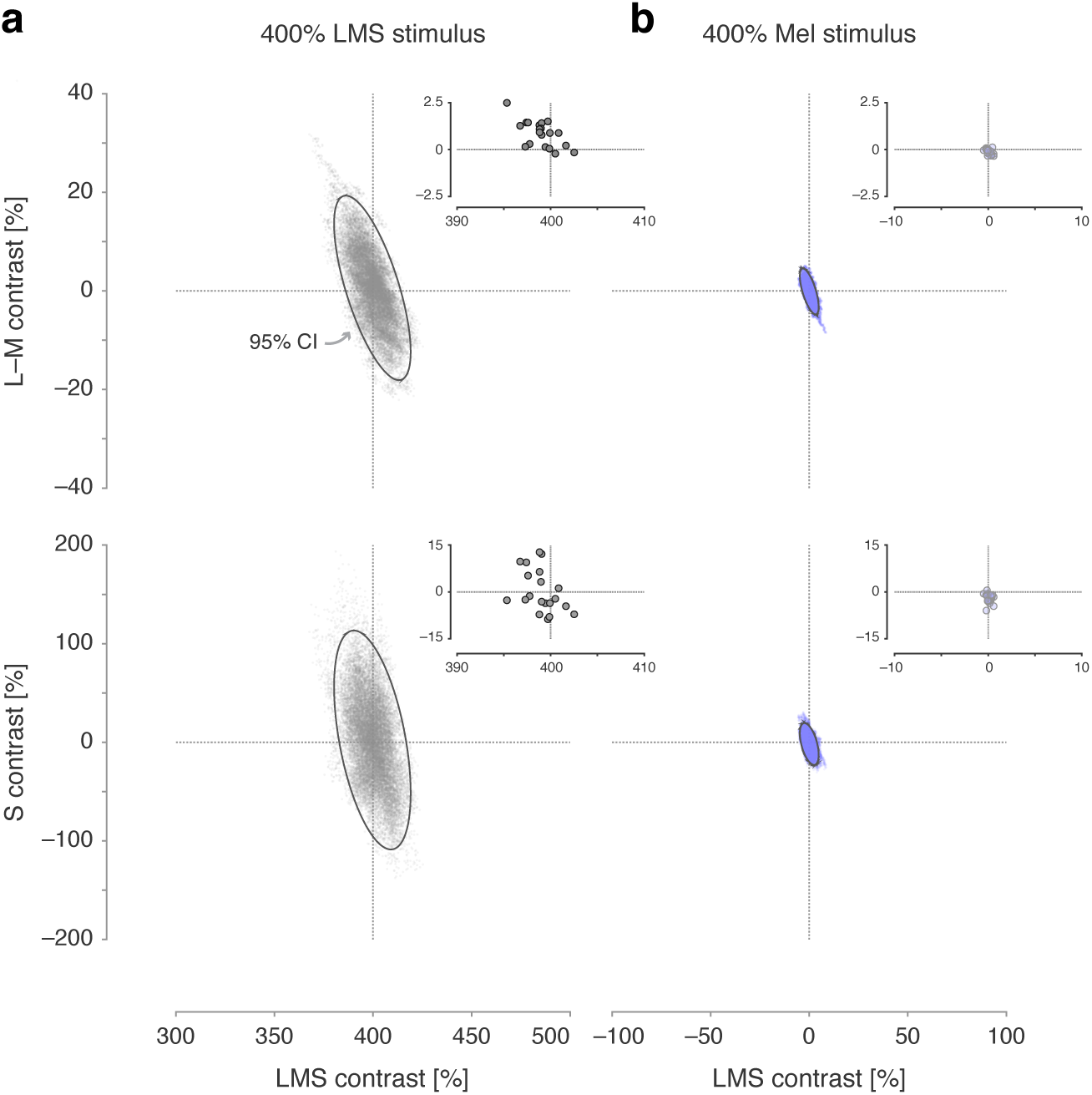
Inadvertent cone contrast in the perceptual stimuli (related to Figure 5). *Inset* in each plot is the calculated post-receptoral cone contrast of the melanopsin and luminance 400% spectral pulses used in the perceptual experiment. Each point corresponds to the difference between the background and stimulus spectra measured for each subject at the time of their testing session. Following the same procedure as described in Figure S5, we then simulated the post-receptoral cone contrast that might be produced by our stimuli in the face of biological variability in our subjects. **(a)** Post-receptoral contrast estimated from simulations for the 400% LMS (luminance) stimulus. **(b)** Post-receptoral contrast estimated from simulations for the 400% melanopsin stimulus.

**Table S1:** Across-subject ratings of nine perceptual qualities for 400% contrast pulses of the three stimulus types.

| *quality* | *melanopsin*<br>median ± inter-quartile range | *LMS*<br>median ± inter-quartile range | *Light flux*<br>median ± inter-quartile range |
| --- | --- | --- | --- |
| cool to warm | 4.25 ± 4.00 | 4.50 ± 2.00 | 4.50 ± 1.25 |
| dull to glowing | 4.50 ± 3.75 | 5.00 ± 1.25 | 5.50 ± 2.50 |
| colorless to colored | 5.75 ± 1.25 | 3.25 ± 2.25 | 2.00 ± 1.50 |
| focused to blurred | 5.00 ± 1.75 | 3.50 ± 3.00 | 3.50 ± 2.25 |
| slow to rapid | 4.00 ± 2.50 | 4.75 ± 2.00 | 4.50 ± 1.75 |
| pleasant to unpleasant | 4.75 ± 2.75 | 3.00 ± 2.00 | 3.00 ± 1.50 |
| dim to bright | 2.50 ± 3.25 | 5.25 ± 1.50 | 6.00 ± 1.25 |
| smooth to jagged | 3.50 ± 2.75 | 2.25 ± 1.75 | 2.00 ± 1.50 |
| constant to fading | 5.00 ± 2.50 | 2.00 ± 1.75 | 2.00 ± 1.50 |

**Table S2:**
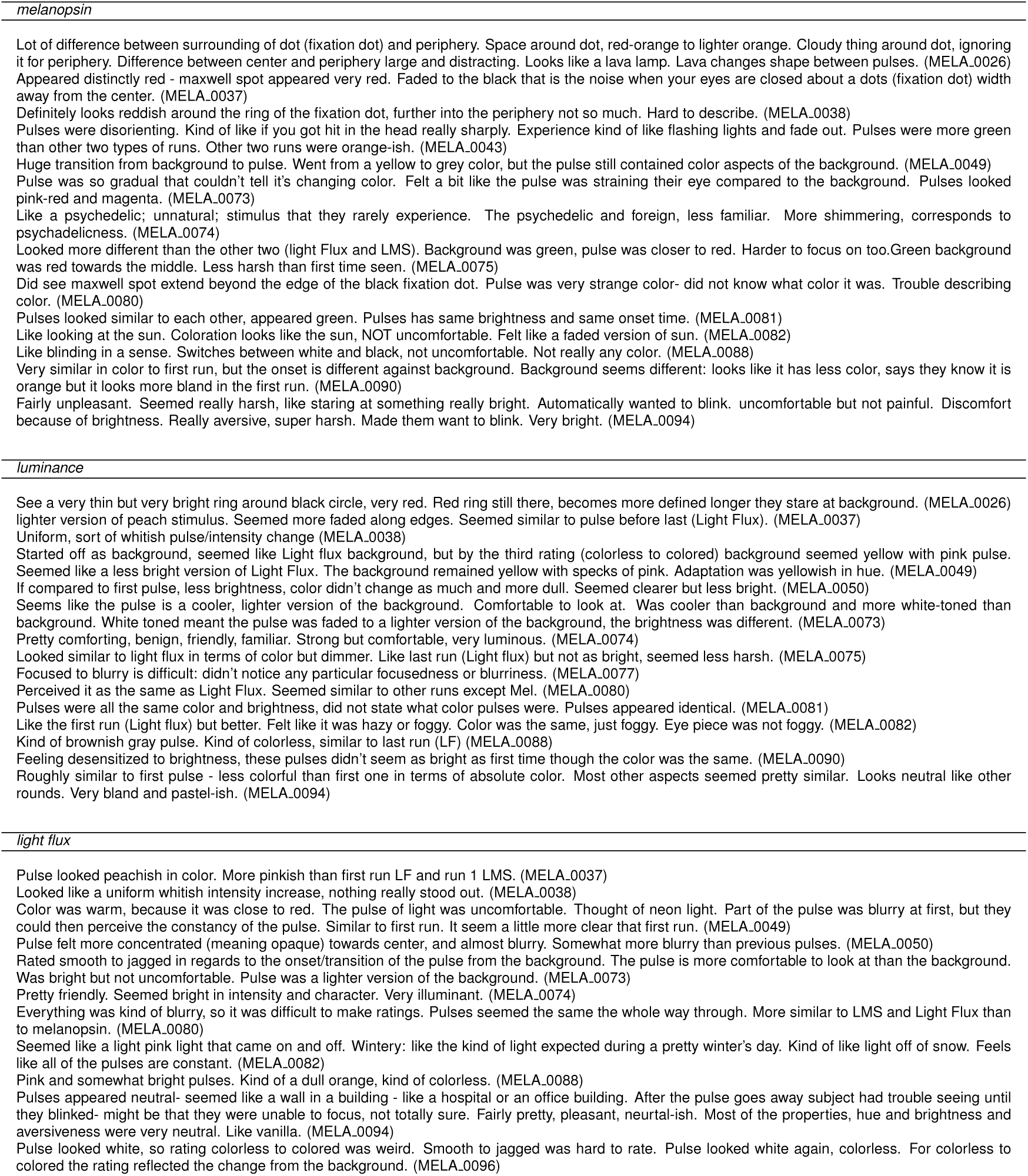
Free-form descriptions of the pulsed stimuli. Subjects in the perception experiment were invited to describe their impressions of the stimuli during a debriefing session and these were recorded by the examiner. Subject ID codes given in parentheses. The subject was not told the spectral identity of the stimuli, and in their descriptions referred to the stimuli by their experimental order; these references to run order are replaced here by the spectral identity of the stimulus for clarity. Some subjects provided descriptions of changes in the appearance of the stimulus at the edge of the masked macular region; they were asked to ignore this aspect of the stimulus in their ratings.

**Table S3:**
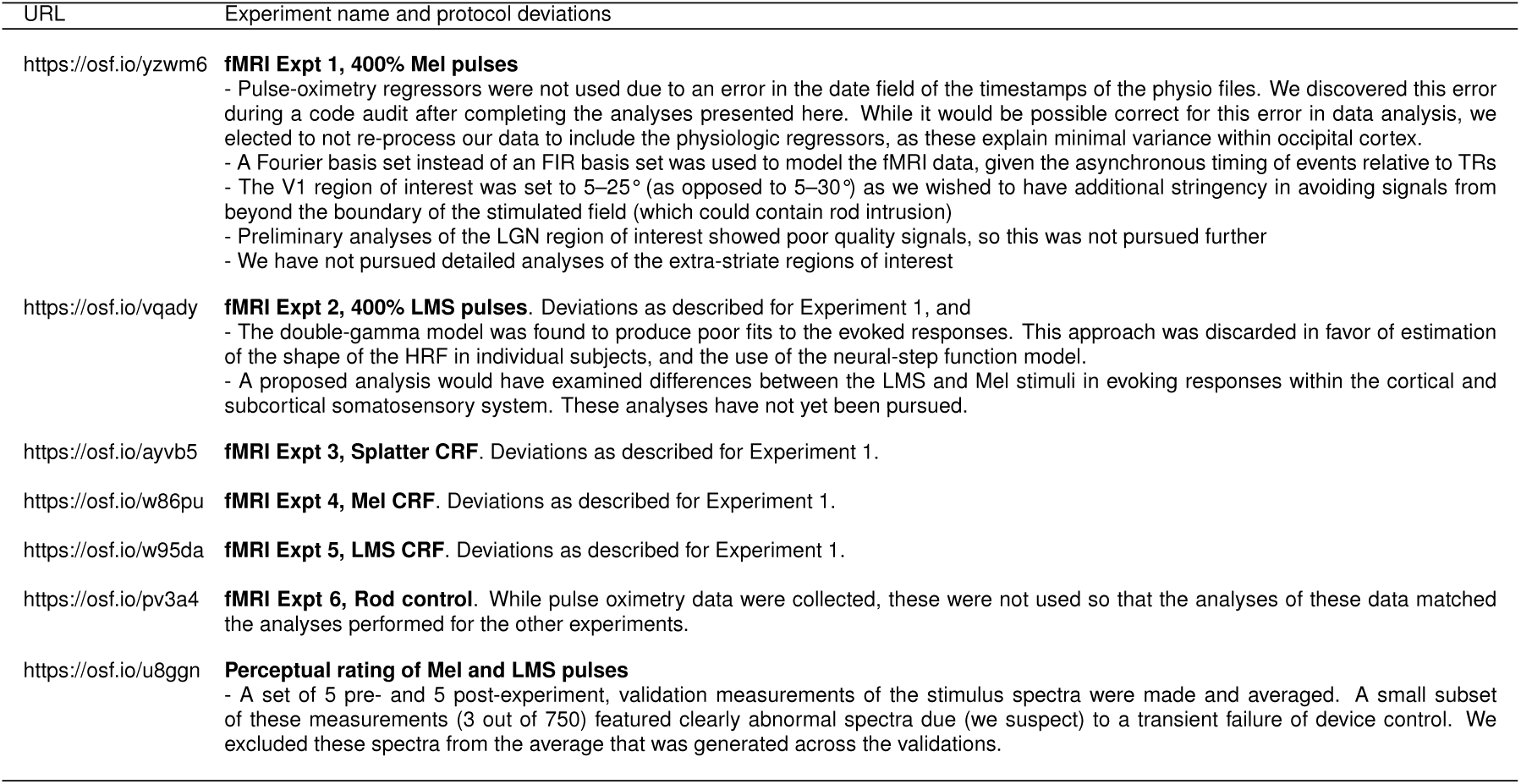
Pre-registrations and protocol deviations. Links are to pre-registration pages on the Open Science Framework site. Some pre-registrations include addenda.

| URL | Experiment name and protocol deviations |
| --- | --- |
| https://osf.io/yzwm6 | **fMRI Expt 1, 400% Mel pulses**<br>- Pulse-oximetry regressors were not used due to an error in the date field of the timestamps of the physio files. We discovered this error during a code audit after completing the analyses presented here. While it would be possible correct for this error in data analysis, we elected to not re-process our data to include the physiologic regressors, as these explain minimal variance within occipital cortex.<br>- A Fourier basis set instead of an FIR basis set was used to model the fMRI data, given the asynchronous timing of events relative to TRs<br>- The V1 region of interest was set to 5–25° (as opposed to 5–30°) as we wished to have additional stringency in avoiding signals from beyond the boundary of the stimulated field (which could contain rod intrusion)<br>- Preliminary analyses of the LGN region of interest showed poor quality signals, so this was not pursued further<br>- We have not pursued detailed analyses of the extra-striate regions of interest |
| https://osf.io/vqady | **fMRI Expt 2, 400% LMS pulses.** Deviations as described for Experiment 1, and<br>- The double-gamma model was found to produce poor fits to the evoked responses. This approach was discarded in favor of estimation of the shape of the HRF in individual subjects, and the use of the neural-step function model.<br>- A proposed analysis would have examined differences between the LMS and Mel stimuli in evoking responses within the cortical and subcortical somatosensory system. These analyses have not yet been pursued. |
| https://osf.io/ayvb5 | **fMRI Expt 3, Splatter CRF.** Deviations as described for Experiment 1. |
| https://osf.io/w86pu | **fMRI Expt 4, Mel CRF.** Deviations as described for Experiment 1. |
| https://osf.io/w95da | **fMRI Expt 5, LMS CRF.** Deviations as described for Experiment 1. |
| https://osf.io/pv3a4 | **fMRI Expt 6, Rod control.** While pulse oximetry data were collected, these were not used so that the analyses of these data matched the analyses performed for the other experiments. |
| https://osf.io/u8ggn | **Perceptual rating of Mel and LMS pulses**<br>- A set of 5 pre- and 5 post-experiment, validation measurements of the stimulus spectra were made and averaged. A small subset of these measurements (3 out of 750) featured clearly abnormal spectra due (we suspect) to a transient failure of device control. We excluded these spectra from the average that was generated across the validations. |

